# Gigavalent display of proteins on monodisperse polyacrylamide hydrogels as a versatile modular platform for functional assays and protein engineering

**DOI:** 10.1101/2021.10.30.466587

**Authors:** Thomas Fryer, Joel David Rogers, Christopher Mellor, Ralph Minter, Florian Hollfelder

## Abstract

The robust modularity of biological components that are assembled into complex functional systems is central to synthetic biology. Here we apply modular “plug and play” design principles to a microscale solid phase protein display system that enables protein purification and functional assays for biotherapeutics. Specifically, we capture protein molecules from cell lysates on polyacrylamide hydrogel display beads (‘PHD beads’), made in microfluidic droplet generators. These monodisperse PHD beads are decorated with predefined amounts of anchors, methacrylate-PEG-benzylguanine (BG) and methacrylate-PEG-chloroalkane (CA). Anchors form covalent bonds with fusion proteins bearing cognate tag recognition (SNAP and Halo-tags) in specific, orthogonal and stable fashion. Given that these anchors are copolymerised throughout the 3D structure of the beads, proteins are also distributed across the entire bead sphere, allowing attachment of ∼10^9^ protein molecules per bead (Ø 20 *μ*m). This mode of attachment reaches a higher density than possible on widely used surface-modified beads, and additionally mitigates surface effects that often complicate studies with proteins on beads. We showcase a diverse array of protein modules that enable the secondary capture of proteins, either non-covalently (IgG and SUMO-tag) or covalently (SpyCatcher, SpyTag, SnpCatcher and SnpTag). Proteins can be displayed in their monomeric forms, but also reformatted as a multivalent display (using secondary capture modules that create branches) to test the contributions of avidity and multivalency towards protein function. Finally, controlled release of modules by irradiation of light is achieved by incorporating the photocleavable protein PhoCl: irradiation severs the displayed protein from the solid support, so that functional assays can be carried out in solution. As a demonstration of the utility of valency engineering, an antibody drug screen is performed, in which an anti-TRAIL-R1 scFv protein is released into solution as monomers-hexamers, showing a ∼50-fold enhanced potency in the pentavalent format. The ease of protein purification on solid support, quantitative control over presentation and release of proteins and choice of valency make this experimental format a versatile, modular platform for large scale functional analysis of proteins, in bioassays of protein-protein interactions, enzymatic catalysis and bacteriolysis.

**Table of Contents Graphics:** 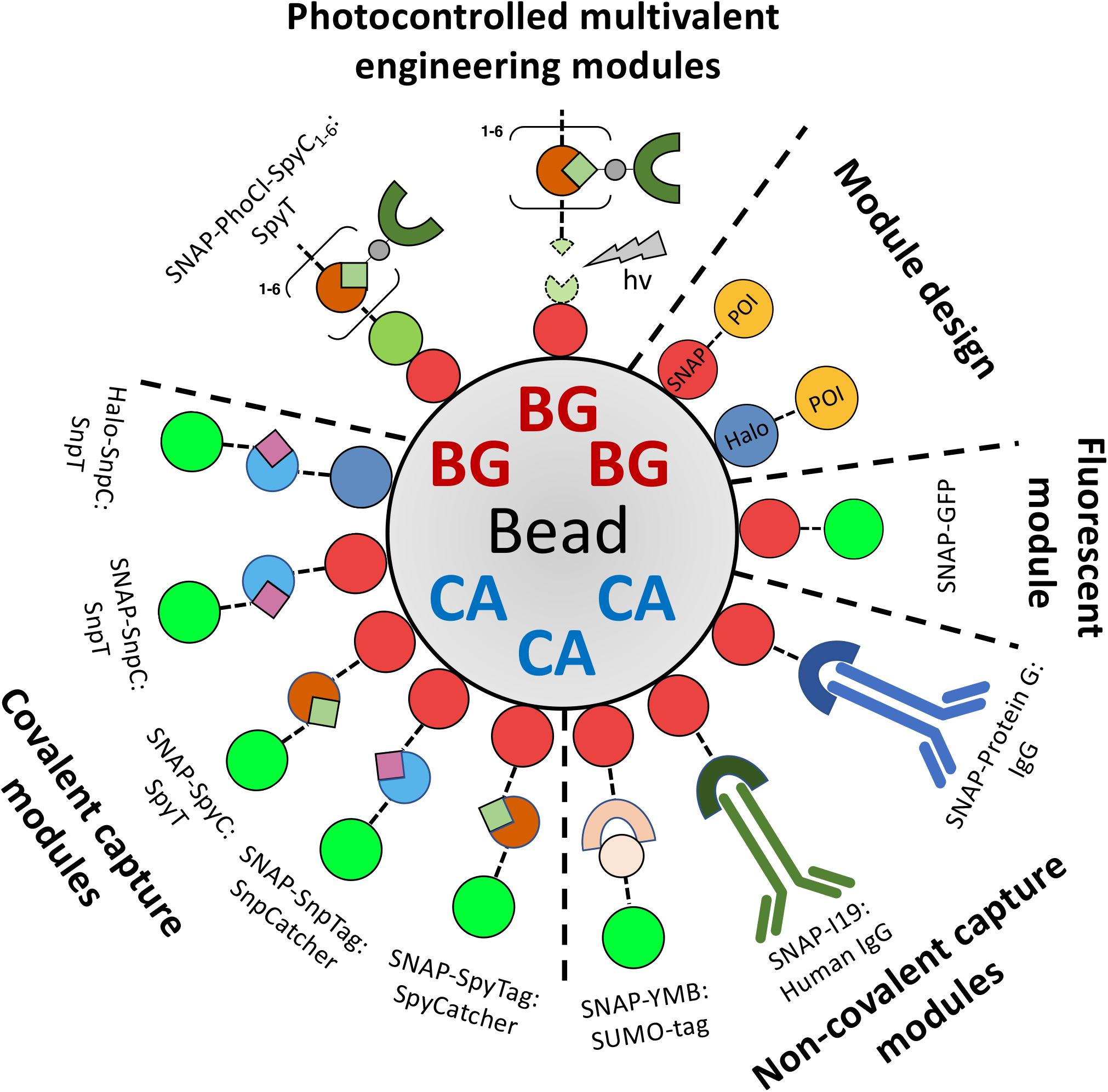

## Introduction

The analysis of proteins, and their use as therapeutics,^1^ enzymes in biocatalysis^2^ and bioremediation,^3^ growth factors for tissue culture^4^ or targets for binder discovery campaigns^5^ is often facilitated by the ability to capture, maintain and manipulate proteins on biocompatible surfaces. Protein solid-phase immobilisation is critical to many bioassays (e.g. ELISA^6^ and SPR^7^ for investigating protein:protein interactions) as it enables washing, modification or rebuffering steps and interfaces with robotic workflows, using the protein attachment to handle the protein for testing in assays or for direct analyses. Industrial-scale biocatalysis can be enhanced by the sequestration/immobilisation of valuable enzymes^2^ in continuous flow biocatalysis,^8,9^ whilst also offering potential synergistic effects through the co-localisation of specific enzymes.^10^ Proteins immobilised on surfaces have also emerged as useful therapeutic agents, enhancing *in vivo* half-life and providing extra control over drug delivery (both temporally and spatially)^1,11^. Despite the demonstrated utility of immobilised proteins across multiple fields, the methods of immobilisation are highly diverse and typically bespoke. In addition, protein function and stability can be impacted by surface effects (observed e.g. for immobilised targets in phage display^5,12^ and enzymes in biocatalysis^13–15^); spectroscopic interference (such as autofluorescence^16^) can negatively affect bioassay sensitivity; the stoichiometry and strength of attachment is variable on heterogeneous solid-phase supports; and there can be batch-to-batch variation that hampers the development of robust and reproducible protocols (‘beads kill leads’).

To address these challenges, new technologies to facilitate assay development and accelerate drug discovery pipelines are needed, based on readily synthesised reagents that can be combined as modules to broaden applicability in robust, reproducible procedures. The importance of versatility for such reagents can be conceptualised by analogy to the field of computer programming, in which there is considerable ongoing effort to produce high-quality, open-source code, and to build accessible user-facing programs based on such code: the goal is that most users’ needs will be satisfied by the package with little or no customisation, but “users” (i.e. researchers) are also able to understand, extend and personalise the program to perform more specialised tasks. The two design principles of modularity and robustness are thus of great importance to make a protein immobilisation system widely applicable. Robustness can be seen as the stability of protein capture over time and under different conditions, as well as the simplicity of the initial capture methodology. Robust protein capture through the trusted modular assembly of “plug and play” components thus provides molecular Lego that simplifies the design of e.g. synthetic biology^17^ experiments, just as click chemistry^18,19^ has made aspects of synthetic chemistry generalisable, easy to use and versatile. The interface of protein engineering and synthetic biology has produced a rich vein of technologies in recent years, notably: the development of SpyCatcher, amongst others^20,21^, as a plug and play tool for post-translational valency engineering and protein purification^22^; the development of photocontrollable proteins such as PhoCl^23^ for the spatiotemporal control of protein release *via* light-induced protein backbone cleavage^24^; or new highly stable and versatile protein recognition elements such as the ALFA-tag system^25^. However few, if any, protein immobilisation methods (e.g. Ni-NTA, Streptavidin, Protein A/G, chemical cross-linking) interface with such technologies in a robust, modular and highly controlled manner, severely curtailing the engineerability of protein-based systems using these common technologies at their core. An “open-source” platform based on synthetic biology principles and programmable at the level of DNA sequence would shift the limit of engineerability from the availability of toolsets to the end-user’s creativity.

Of the surfaces functionalised with proteins, hydrogels^26^ are an increasingly important matrix for biological applications due to their biocompatibility (permeability, adjustable stiffness, low cytotoxicity). They have found use in single-cell transcriptomics^27^, mammalian cell culture^28^, *in vivo* drug delivery devices^1^ or as artificial cells^29^. In particular, surface effects can be minimised by the absence of a hydrophobic surface that can lead to protein denaturation. Hydrogels functionalised with protein have been demonstrated utilising a diverse array of capture methods (anti-His-tag aptamers^29^, molecular imprinting^30^, click chemistry^24^, and co-polymerisation with acrylamide^31^ or through disulphide bond formation^32^), yet no simple, modular, and site-specific method has been developed.

Here, we introduce a platform that incorporates covalent, site-specific protein capture in highly modular fashion that offers stability, versatility, and accessibility. Using this technology suite, proteins can be captured at precisely defined valencies, in a highly specific and orthogonal manner, and released on demand by exposure to light, so that high-throughput affinity and enzymatic assays, bioassay sensor designs and molecular engineering strategies of biologics can be implemented.

## Results and Discussion

### Design of polyacrylamide hydrogels with titratable protein capture

Synthesised from components found in most molecular biology labs (e.g. to make SDS-PAGE gels) and with a proven reliability of polymerisation, polyacrylamide hydrogels are easy to use and have already taken a role as biocompatible scaffolds for the delivery of reagents in microfluidic single-cell transcriptomic workflows.^27^ However no simple, stable, modular technology exists for the functionalisation of polyacrylamide hydrogels with proteins. Polyacrylamide hydrogels consist of chains of monomers of acrylamide cross-linked by bis-acrylamide in stable polymers and thus, to enable the capture of proteins, we copolymerised acrylamide and bis-acrylamide monomers with methacrylate-modified small molecule ligands (methacrylate-PEG-benzylguanine (BG) and -chloroalkane (CA); ***Figure 1a***). These ligands act as suicide substrates for SNAP-tag^33^ and Halo-tag^34^ respectively, and their copolymerisation throughout the hydrogel enables completely covalent capture of an array of modular building blocks expressed as fusion proteins to these tags (***Figure 1b***). SNAP-tag and Halo-tag are both well-established protein tags, used across biological fields and can be expressed in bacterial, yeast and mammalian cell lines^33^. Notably SNAP-tag and Halo-tag react entirely orthogonally with their respective ligands (BG and CA), and have already been used to capture proteins on surfaces^35,36^, yet this orthogonality has not been fully exploited for protein capture on bifunctional surfaces, and “plug and play” modules for protein engineering and assay design have not been developed.

**Figure 1.**
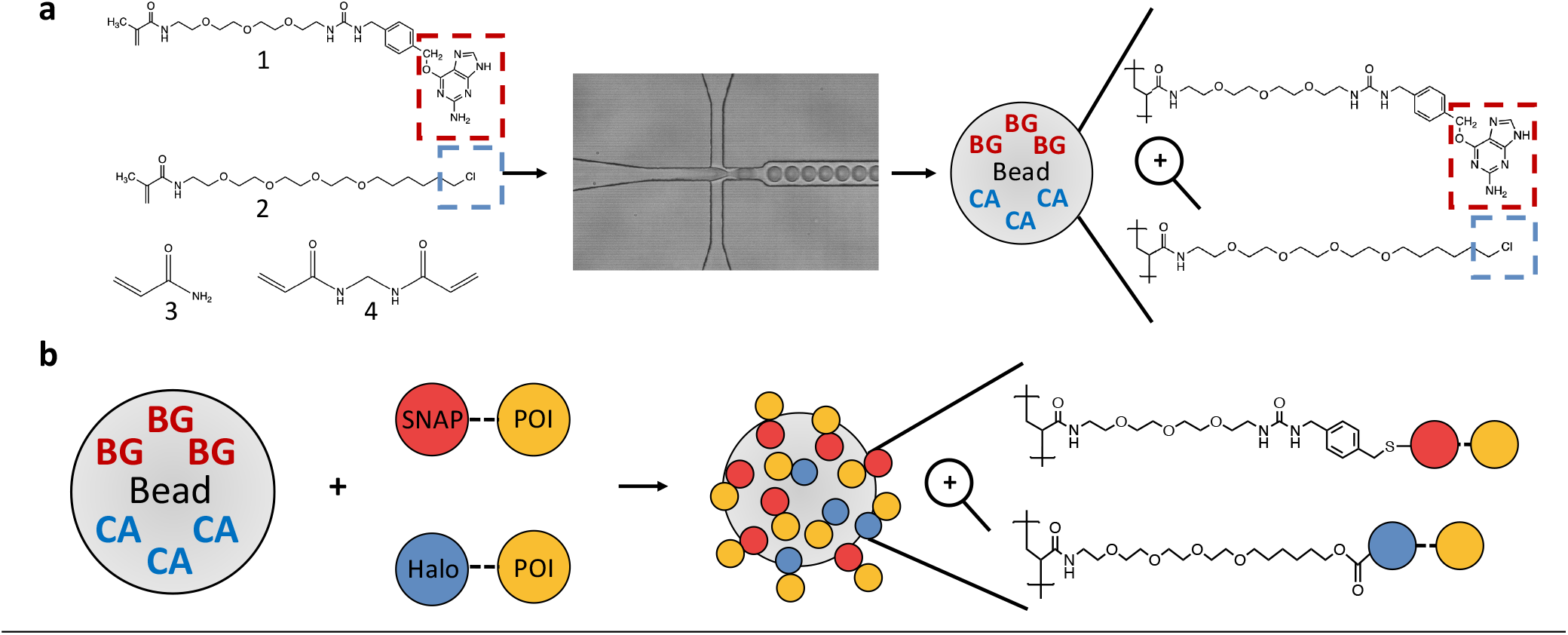
Modular polyacrylamide hydrogel display (**a**) Monodisperse polyacrylamide hydrogel beads are made through the encapsulation of monomers (1: Methacrylate-PEG-*b*enzyl*g*uanine (BG), 2: Methacrylate-PEG-*c*hloro*a*lkane (CA), 3: Acrylamide, 4: Bis-acrylamide) with polymerisation-inducing catalysts using droplet-based microfluidics. Upon de-emulsification BG (*red*) and/or CA (*blue*) are retained within each bead due to co-polymerisation with the hydrogel backbone (**b)** Hydrogel beads can then be orthogonally functionalised with SNAP- or Halo-tag fusion proteins (red and blue, respectively) through covalent reaction with their respective co-polymerised small molecule ligands (BG/CA).

We prepared methacrylate-PEG-benzylguanine/-chloroalkane by reacting methacrylate-NHS ester with amine-PEG-benzylguanine or -chloroalkane overnight in a simple click reaction and achieved near-quantitative yield (>90 %, as measured by HPLC, ***Figure S1*.*1, Table S1*.*1***). The products of these reactions can then be directly used for co-polymerisation into polyacrylamide hydrogels, and so we subsequently generated BG-functionalised monodisperse beads of 20 *μ*m diameter (Ø) using droplet-based microfluidics at ∼8 kHz (enabling production of 29 million beads per hour). 20 *μ*m beads are readily compatible with downstream analysis technologies such as flow cytometry. However, any desired size can be made through the use of different chip geometries and flow rates. Upon de-emulsification, BG-functionalised hydrogel beads can be incubated with SNAP-tag fusion proteins (such as SNAP-GFP) for covalent capture (***Figure 2a)***. Specific protein capture is exemplified in ***Figure 2b***: only beads functionalised with BG are able to capture SNAP-GFP, and there is little to no non-specific binding to non-functionalised polyacrylamide beads. PHD beads can also be made entirely *without* the use of microfluidics, by vortexing the aqueous monomer solution with surfactant-containing oil (the same compositions as for microfluidics) to create polydisperse emulsions. These polydisperse hydrogel beads vary somewhat in size but are still highly functional for capture of e.g. SNAP-GFP (***Figure S1.2***), and so this technology is also accessible to researchers without a microfluidic set-up.

**Figure 2.**
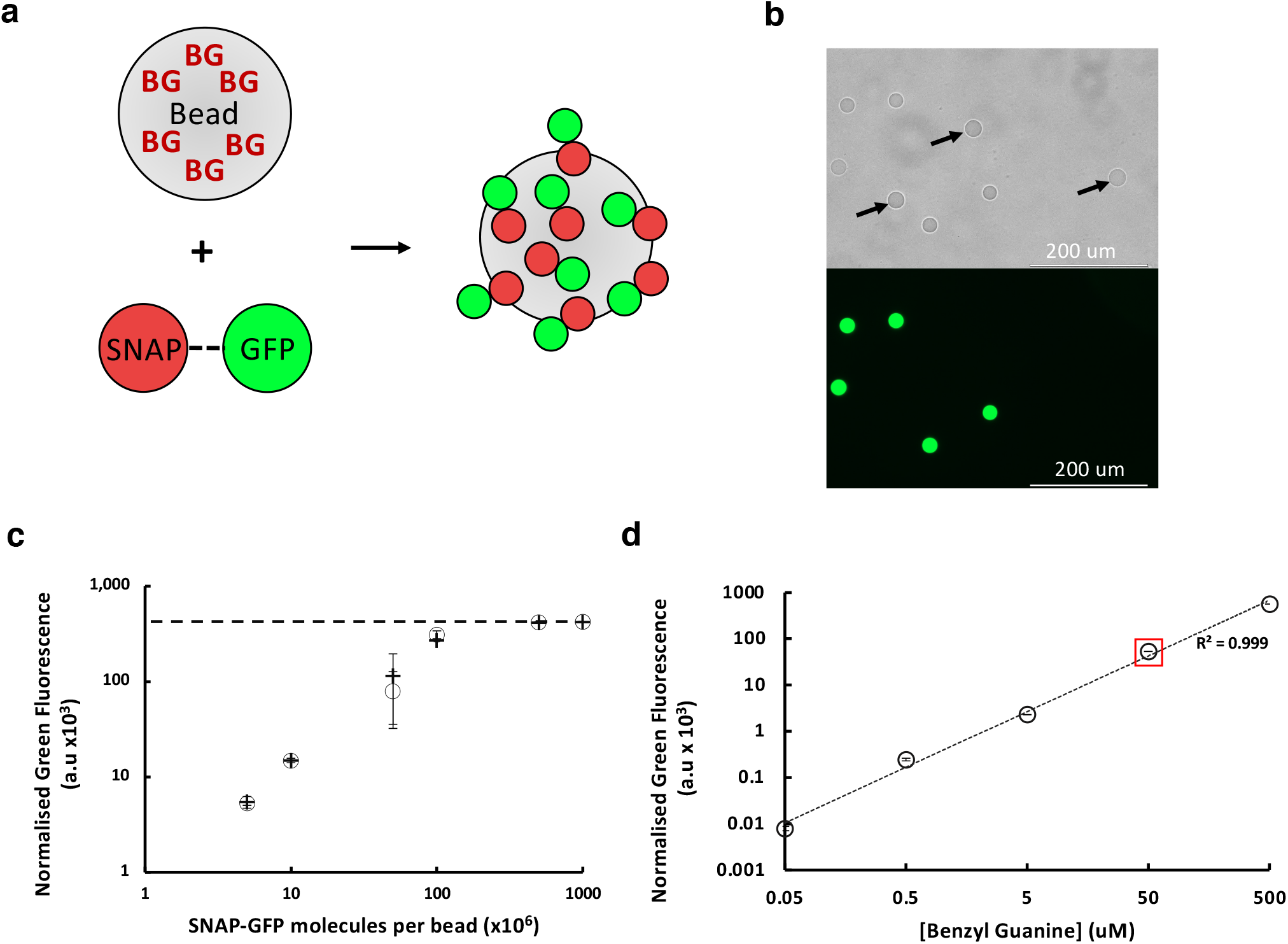
Specific, stable and titratable protein capture on polyacrylamide hydrogel beads. (**a**) BG functionalised hydrogel beads are incubated with SNAP-GFP, leading to the covalent capture of SNAP-GFP on bead. (**b**) 20 *μ*m PHD beads +/- 50 μM BG were mixed 50:50 and incubated with SNAP-GFP followed by washing and imaging (top panel brightfield, bottom panel GFP channel) to detect specific GFP attachment. Scalebar: 200 μm. Arrows indicate negative beads (**c**) 100,000 of 20 *μ*m, 50 *μ*M BG PHD beads were incubated with defined numbers of SNAP-GFP molecules per bead overnight, washed and analysed by flow cytometry. The saturation point, i.e. where the addition of extra SNAP-GFP does not lead to an increase in on-bead fluorescent signal (dashed line) corresponds to a density of ∼150 million attached proteins per bead. Black crosses indicate boiled beads, open circles indicate beads handled according to our standard procedure (see Methods section). (**d**) Five sets of 20 *μ*m PHD beads were prepared with the indicated BG loading. All were incubated with an excess of SNAP-GFP, washed, and analysed by flow cytometry. The red square highlights the 50 μM BG beads used in (c), that captured 1.5 × 10^8^ SNAP-GFP molecules per bead, the near-perfect correlation between [BG] and green fluorescence shows that a valency range of 10^5^-10^9^ per bead was achieved (for 0.05, 0.5, 5, 50 and 500 μM BG beads respectively). Data are the mean of triplicates, normalised to the background signal of PHD beads lacking BG.

Next, we sought to quantify the capacity of on-bead coupling. When incubating beads (Ø 20 *μ*m, 50 *μ*M BG) with increasing amounts of SNAP-GFP, we observed asymptotic saturation of the fluorescence signal (after washing of beads) at approximately 5 × 10^8^ SNAP-GFP molecules per bead, and we found this binding behaviour to be highly conserved even when beads are boiled before protein capture, demonstrating the high stability of this system (***Figure 2c***). In order to estimate, more accurately, the number of molecules required to saturate a bead, we extrapolated the linear part of our saturation curve up to the asymptote. This calculation suggests that ∼1.5 × 10^8^ SNAP-GFP molecules per bead are bound at saturation (equal, within experimental error, to the calculated 1.3 × 10^8^ BG molecules per (Ø 20 *μ*m, 50 *μ*M BG) bead; ***Figure S1*.*3a***). Such high occupation levels of immobilised proteins exceed those achieved with magnetic beads that bind proteins on their surface by three orders of magnitude^37^ (M-280 Streptavidin Dynabeads ∼6.6 × 10^5^ IgG molecules per bead ***Figure S1*.*3b***). The difference can be ascribed to the voluminal nature of protein capture, wherein not only is the bead’s surface functionalised, but also its interior. In addition to the high levels of protein capture, it is also possible to precisely control the amount of captured protein by changing the concentration of BG monomers included in the hydrogel polymerisation mix. When the concentration of BG in the initial one-pot pre-polymerisation acrylamide mix is varied, the amount of SNAP-GFP captured varies correspondingly; display densities spanning at least five orders of magnitude can be brought about at will, and an estimated 1.5 × 10^9^ molecules are bound when using 500 uM BG (***Figure 2d***), demonstrating gigavalent capture.

### Specific protein capture via non-covalent secondary capture modules

To overcome the need for *ad hoc* solutions in assay design, we next developed a modular design principle based on the stability and specificity of covalent protein capture on bead. Such a “plug and play” engineering approach enables a researcher to assemble desired protein constructs and apply them to a designed assay or application simply by combining specific modules of defined functions. Whilst we have already demonstrated the direct capture of a protein of interest as a SNAP-tag fusion (***Figure 2***), it is also possible – and greatly enhances the utility of the PHD technology as an engineering tool – to use specific secondary capture modules (e.g. affinity reagents fused to SNAP-tag) to assemble proteins of interest on bead (***Figure 3a***). As the base bead remains the same (20 *μ*m, 50 *μ*M BG), its desired functionality can be altered simply by choosing which secondary capture module to initially capture on bead. To demonstrate this principle, we fused several secondary capture modules to SNAP-tag for immobilisation: SNAP-Protein G for mouse IgG capture (***Figure 3b***); SNAP-I19^38^, an anti-human IgG DARPin, for human IgG capture (***Figure 3c***); and SNAP-YMB^39^, an anti-SUMO monobody, for capture of SUMO-GFP (***Figure 3d***). These secondary capture modules are all readily expressed in bacteria, obviating the need to buy expensive affinity reagents for desired applications. Further secondary capture modules can be designed based on published sequences of affinity reagents or freshly developed through *de novo* discovery techniques such as phage display, and assembled in a modular fashion using e.g. Gibson Assembly (details in ***Figure S1*.*5, Experimental section S2*.*3***).

**Figure 3.**
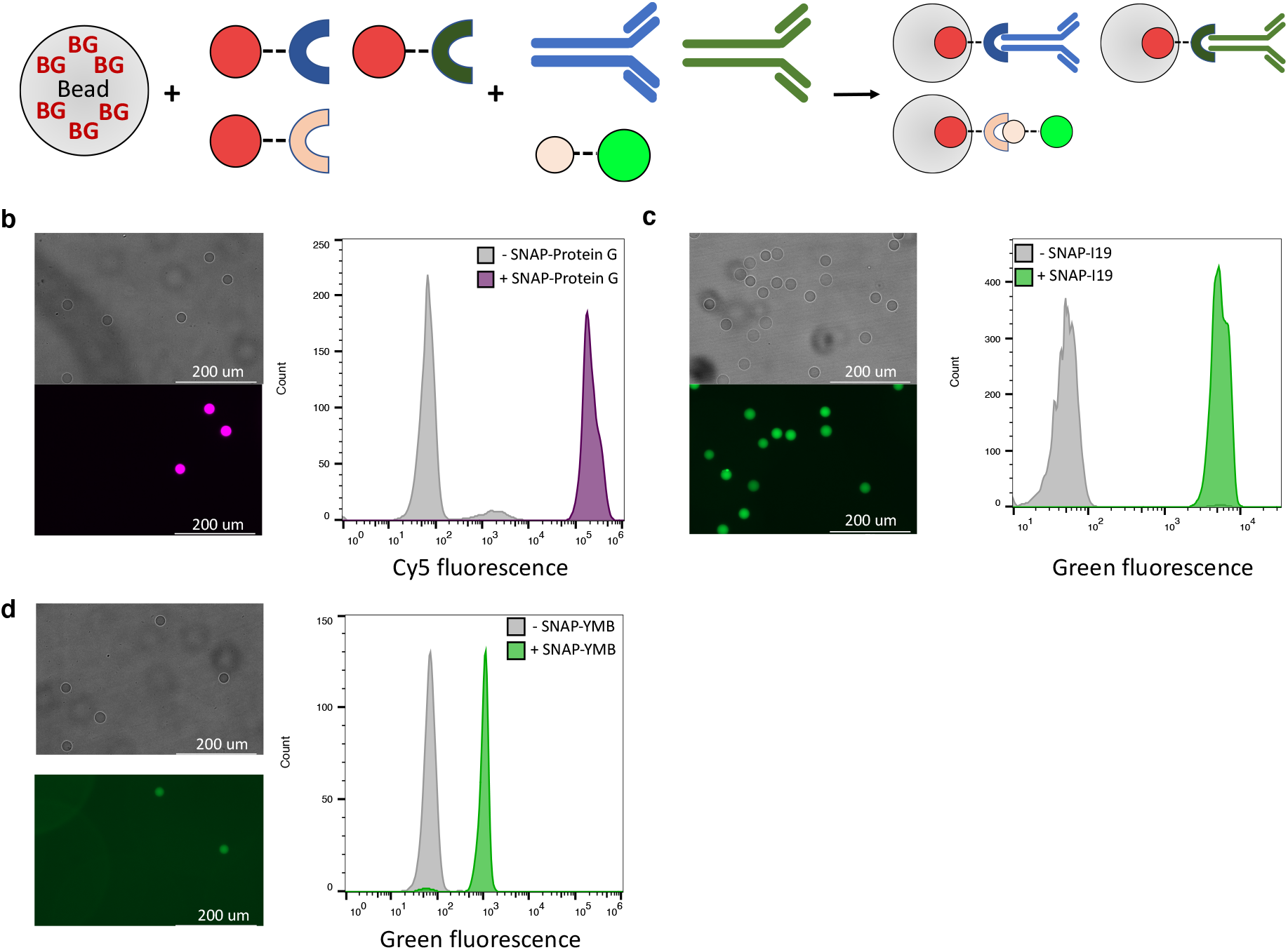
Versatile capture of modular building blocks for specific protein capture. (**a**) BG-functionalised hydrogel beads can be used to covalently immobilise secondary capture modules (as SNAP-tag fusions) that are specific for a desired target protein. (**b-d**) Functionalised PHD beads (Ø20 *μ*m; 50 uM BG) +/- (**b**) SNAP-Protein G, (**c**) SNAP-I19 or (**d**) SNAP-YMB were incubated with their respective target proteins (Mouse IgG-iFluor 647, human IgG1-AlexaFluor 488, SUMO tag-GFP) for 1 hour, washed and analysed by both fluorescent microscopy (left-hand panels, +/- SNAP-tag capture module mixed and imaged together) and flow cytometry (right-hand panels, +/- SNAP-tag capture module analysed separately and super-imposed). Scale bars represent 200 *μ*m.

### Modular and orthogonal programming of bead functionality via covalent secondary capture modules

Next, to enable *covalent* immobilisation of proteins of interest, we designed additional secondary capture modules as both SNAP- and Halo-tag fusions to the suite of SpyCatcher/SpyTag and SnpCatcher/SnpTag technologies^40^ (***Figure 4a***). These protein pairs form an isopeptide bond under standard biological reaction conditions and have already been applied widely to the modular engineering of proteins (e.g. vaccine design^41^, protein cyclisation for enzyme engineering^42^, multivalent and multifunctional protein assembly^43^). In this work we use SpyCatcher ΔNC^44^ (a deimmunised SpyCatcher truncation) and SpyTag002^45^ (an evolved SpyTag with enhanced reaction kinetics). Importantly the two pairs react orthogonally (as do SNAP-tag and Halo-tag) enabling the specific modular construction of multifunctional beads with relative ease. Whilst SpyTag and SnpTag have both been incorporated into hydrogel frameworks previously (PEG-functionalised^31^, or all-protein hydrogels^46^) the versatility of these systems is limited compared to that displayed here in which any and all arrangements of protein pairs can be assembled on bead (***Figure 4 b-e***) simply by exchanging the covalent secondary capture module first captured on bead. Due to the orthogonality of the four protein capture technologies employed (SNAP-tag, Halo-tag, SpyCatcher, SnpCatcher), specific capture of target proteins can be programmed by simply functionalising beads +/- any desired component. In ***Figure 4f*** we demonstrate the highly controlled capture of GFP-SnpT/mCherry-SpyT based solely upon the previous functionalisation of beads with/without SNAP-SpyCatcher/Halo-SnpCatcher. These beads now exhibit programmed bifunctionality (both GFP and mCherry fluorescence), and serve to demonstrate the versatility, modularity and orthogonality of the PHD technology. As before, researchers can design and express further capture modules and functionalities with relative ease through the use of modular Gibson Assembly.

**Figure 4.**
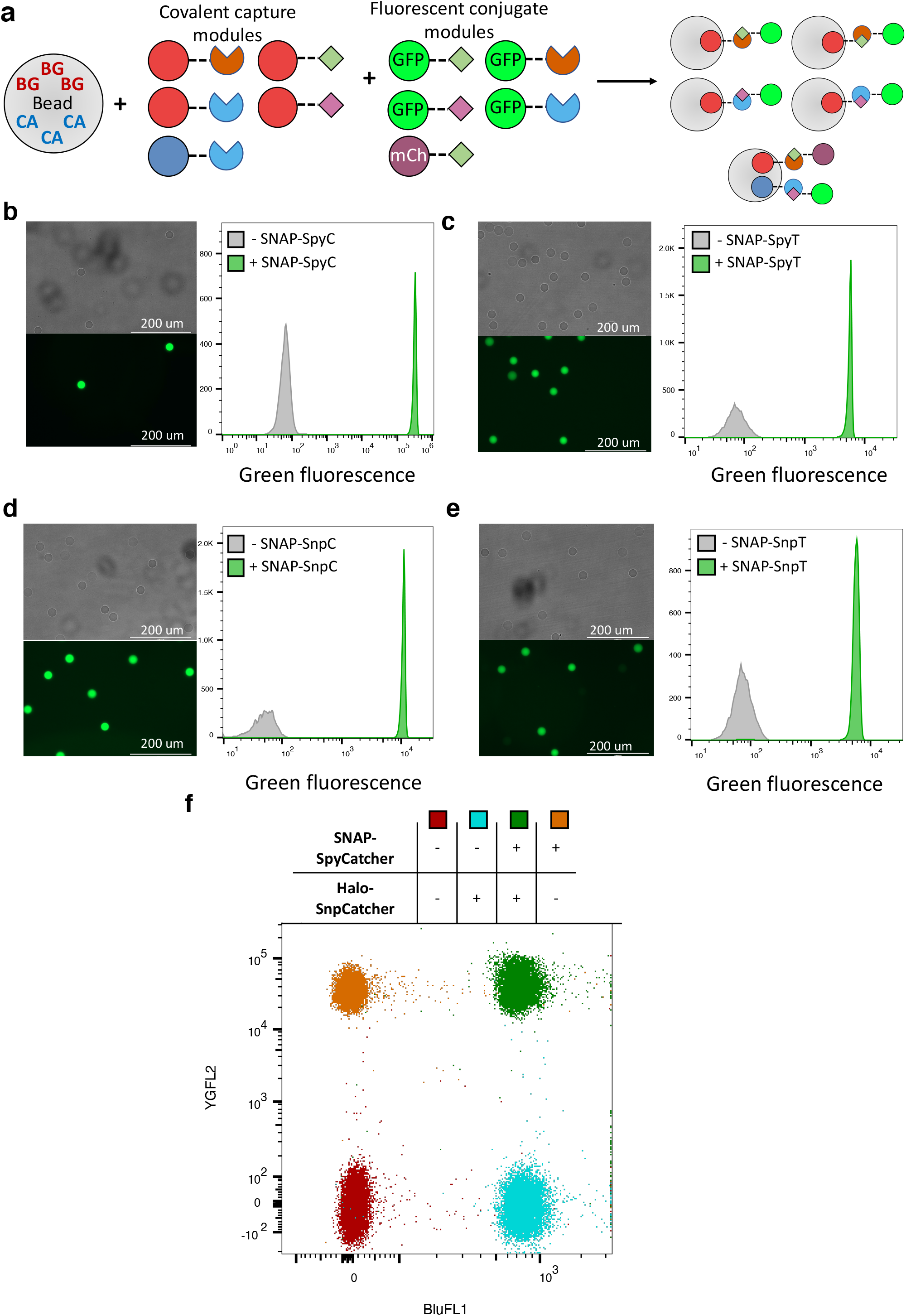
Versatile, orthogonal and covalent capture of target proteins. (**a**) BG/CA-functionalised hydrogel beads can be used to covalently immobilise secondary covalent capture modules (as SNAP- tag or Halo-tag fusions) that specifically react with their partner tag. (**b-e**) Monofunctionalised PHD beads (50 *μ*M BG, Ø 20 *μ*m) were incubated +/- SNAP fusion proteins: **(b)** SNAP-SpyCatcher; (**c**) SNAP-SpyTag; (**d**) SNAP-SnpCatcher; or (**e**) SNAP-SnpTag. These beads were then mixed and incubated with their respective target proteins (*b*: GFP-SpyTag; *c*: GFP-SpyCatcher; *d*: GFP-SnpTag; *e*: GFP-SnpCatcher) for 1 hour, washed and analysed by both fluorescent microscopy (left-hand panels) and flow cytometry (right-hand panels). Scale bars represent 200 *μ*m. (**f**) Bifunctionalised PHD beads (50 *μ*M BG, 50 *μ*M CA, Ø 20 *μ*m) were incubated +/- SNAP-SpyCatcher and/or Halo-SnpCatcher), these beads were subsequently incubated with both GFP-SnpTag and mCherry-SpyTag for 1 hour, washed and analysed by flow cytometry.

### Application of PHD beads to bioassays: protein-protein interactions, enzymatic catalysis and bacteriolysis

Due to the modularity and robustness of the PHD technology it is facile to design and implement bioassays. We demonstrate this for assaying protein:protein interactions – an extremely common bioassay which is key to understanding basic molecular interactions (e.g. in the development of protein-based therapeutics) – by carrying out an investigation into the binding kinetics of the SpyCatcher-SpyTag pair (***Figure 5a***). We incubated SNAP-SpyCatcher functionalised beads with three different concentrations of GFP-SpyTag for up to 60 minutes, before washing away unreacted GFP-SpyTag and measuring the amount of reaction product by flow cytometry – this approach allowed us to readily investigate the effect of GFP-SpyTag concentration on the reaction kinetics.

**Figure 5.**
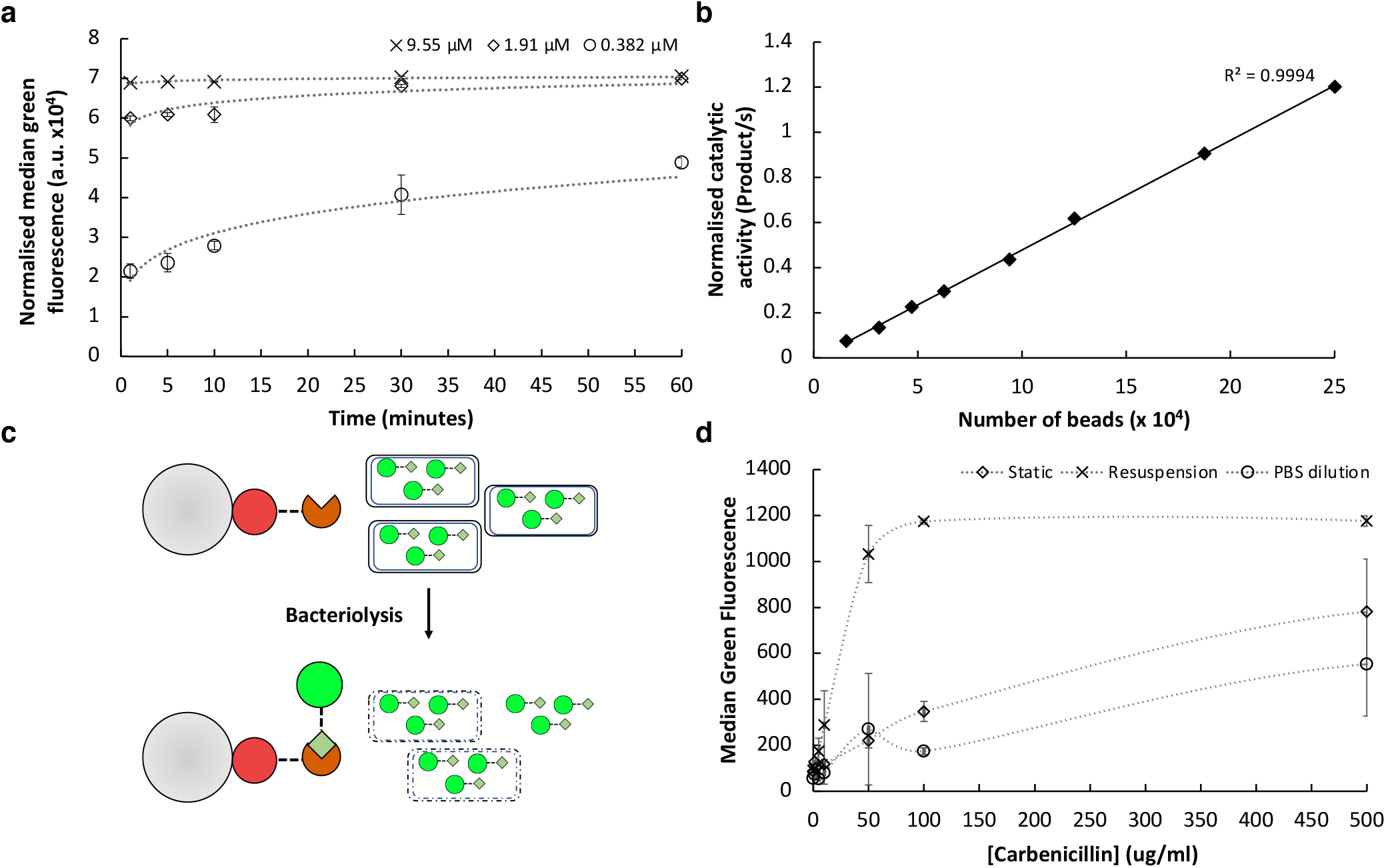
PHD beads in designed functional bioassays. (**a**) PHD beads (50 *μ*M BG, Ø20 *μ*m) functionalised with SNAP-SpyCatcher were incubated with three concentrations of GFP-SpyTag, 9.55 *μ*M (cross), 1.91 *μ*M (open diamond), or 0.382 *μ*M (open circle) at room temperature with rolling for the indicated times. Beads were recovered, washed, and analysed by flow cytometry. Data is presented normalised to non-functionalised beads, and was acquired in duplicate (**b**) P91-SpyTag was captured on bead, washed and incubated with 50 *μ*M substrate (fluoresceine-di(diethylphosphate)) in 100 *μ*l volume. Bead number per well was varied as indicated. The initial 90 minutes of reaction was used to calculate the catalytic activity. Data is presented normalised to non-functionalised beads. (**c**) Overview of bacteriolysis sensor design. PHD beads functionalised with the SNAP-SpyC covalent capture module are incubated with bacterial cells expressing GFP-SpyT. Only upon lysis will the GFP-SpyT be released into solution and be able to be captured on the sensor beads (***d***) *E. coli* cells expressing GFP-SpyTag were grown overnight with induction of protein expression. Static cultures (blue), cultures resuspended in fresh culture media (orange), and static cultures diluted 1:1 with PBS (grey) were incubated with a range of carbenicillin concentrations for 90 minutes at 37 °C in triplicate. Cultures were pelleted and the supernatant transferred to incubate with SpyCatcher-functionalised PHD beads for 60 minutes. Beads were washed twice and then analysed by flow cytometry.

Having demonstrated the utility of PHD beads for assaying protein:protein interactions, we wished to also highlight their suitability for simplifying and improving the quality of high-throughput assays. SNAP-SnpTag was captured directly from bacterial cell lysate and probed by subsequent incubation with GFP-SnpCatcher. A minimal volume of cell lysate corresponding to ∼2ul of culture volume (1/500 of the largest volume tested) was found to already saturate 50,000 20 *μ*m, 50 *μ*M BG beads (***Figure S1*.*4***). Direct capture of a protein of interest from cell lysate obviates the need for a separate purification step, whilst the precise control over protein capture through user-controlled BG concentration and bead number effectively achieves expression level normalisation for a subsequent assay. Protein expression, lysis, on-bead capture and the subsequent assay (flow cytometry) were all carried out in a 96 deep-well plate format; combining the PHD beads with high-throughput, sensitive techniques such as flow cytometry creates a powerful platform with which multiple parameters (e.g. affinity and specificity) can be examined simultaneously, and assays can be multiplexed for even greater throughput^47^.

In addition to protein:protein interactions another common form of bioassay is enzymatic catalysis, in which the accumulation of product or loss of substrate is followed over time. The immobilisation of enzymes is of great interest for industrial biocatalysis^2^ and can also serve to provide a simple method of delivering a defined concentration of protein to a given assay – an important feature when comparing the activity of enzyme variants in a directed evolution experiment for instance. To demonstrate the precise control of enzyme concentration for use in a subsequent bioassay we captured P91^48^-SpyTag, a phosphotriesterase, on SNAP-SpyCatcher-functionalised beads. The number of beads per reaction was varied, and the accumulation of product followed by an increase in fluorescence signal (***Figure 5b***). A near-perfect linear relationship is seen between bead number per reaction and catalytic activity, highlighting the compatibility of PHD beads with enzymatic bioassays. In addition, this experiment also highlights the compatibility of PHD beads with the cell-free expression of proteins, as P91-SpyTag was expressed using PURExpress and directly captured on bead from the *in vitro* expression reaction. Cell-free expression of proteins is now a well-established field^49^ with commercial products available, and can enable the rapid, and (ultra) high-throughput expression even of toxic proteins^50^.

Next, to demonstrate that our platform’s applications are not limited to cell-free bioassays, we designed a microtitre-plate and flow cytometry compatible sensor for bacteriolysis to facilitate the discovery of antibacterials (***Figure 5c***). PHD beads were first functionalised with the SNAP-SpyCatcher covalent capture module before being incubated with *E. coli* which expressed GFP-SpyTag intracellularly and had been exposed to carbenicillin at a range of different concentrations (0-500 *μ*g/mL) and under three different conditions: static culture; culture diluted 1:1 in PBS; and culture resuspended in fresh media (***Figure 5d***). Bacteriolysis is sensed by the release of GFP-SpyTag from lysed bacteria and its subsequent capture on SNAP-SpyC functionalised PHD beads. These sensor beads can then be recovered and quantitatively analysed by flow cytometry. We observed that resuspension of cells in fresh media was necessary for the maximal induction of bacteriolysis, and we further note that these results implicate carbenicillin (and/or related molecules) as an effective protein extraction reagent.

### Valency engineering and photocontrolled release of antibody drugs for phenotypic assays

As an extension to the tools already exhibited, we sought to develop a method of releasing captured proteins into solution upon exposure to a specific cue. Ideally this process would be simple, highly controllable and stable, without the requirement for addition of further reagents. Recent advances have enabled the use of genetically encoded photocontrollable elements for micropatterning^24^ and control of hydrogel stiffness^51^ utilising the photocleavable protein PhoCl^23^. Upon exposure to violet light, PhoCl cleaves its own backbone, thus allowing for the controlled release of attached proteins. Previous attempts to use PhoCl for the controlled release of proteins from hydrogels used click chemistry for immobilisation, which can negatively affect protein functionality through non-site-specific protein capture as well as limiting the engineerability of the system through a lack of orthogonality and easy modularity^24^. As such, we designed and tested a new modular building block, SNAP-PhoCl-SpyCatcher, that would release the SpyCatcher and any associated cargo from the hydrogel (***Figure 6a***). Cleavage in solution was first verified, with significant cleavage seen after just one minute of exposure to light (***Figure 6b***). Due to the transparent nature of the PHD beads, we expected photocleavage to retain comparable efficiency when the SNAP-PhoCl-SpyCatcher modular building block is captured on bead. To test this, beads were functionalised with SNAP-PhoCl-SpyCatcher and exposed to 405 nm light. After light exposure (to prevent any effect of photobleaching) beads were incubated with mCherry-SpyTag to assay for PhoCl cleavage, and hence loss of the SpyCatcher entity from bead. Greater than 83% of protein is released after 5 minutes of exposure to 405 nm light (***Figure 6c***). Improved photocleavage proteins, such as the recently developed PhoCl2,^52^ can be easily incorporated based on the modular design.

**Figure 6:**
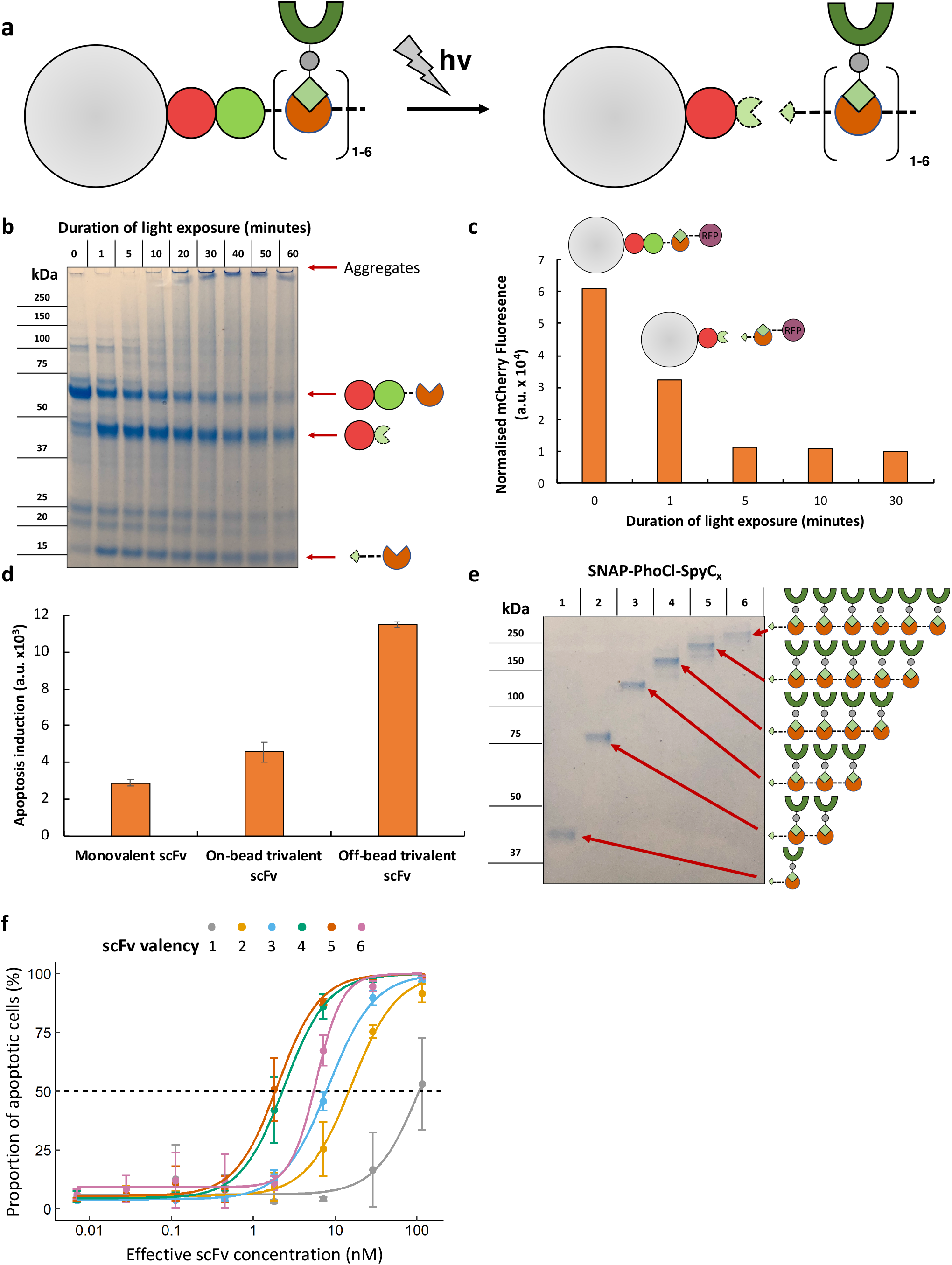
Photocontrolled valency engineering for antibody drug phenotypic assays. (**a**) Beads can be functionalised with the valency engineering covalent capture modules (SNAP-PhoCl-SpyC_1-6_) and subsequently used to capture SpyT-POI (here scFv-SpyT). Upon exposure to 405 nm light the PhoCl protein self-cleaves and releases the valency-modified assembly into solution. (**b**) SNAP-PhoCl-SpyC_1_ was exposed to 405 nm light for the indicated durations and the samples loaded on a denaturing SDS- PAGE gel for analysis of cleavage (**c**) PHD beads functionalised with SNAP-PhoCl-SpyC_1_ were exposed to 405 nm light for the indicated durations. Beads were then washed and incubated with mCherry-SpyTag followed by flow cytometry. Data is presented normalised to PHD beads treated in the same manner, without incubation with mCherry-SpyTag to account for photoswitching of the PhoCl fluorophore upon 405 nm light exposure. (**d**) 100,000 20 *μ*m 50 *μ*M BG beads for each sample were incubated with SNAP-PhoCl-SpyC_3_ and then 3B04-SpyTag. Samples were then treated +/- light and incubated with HeLa cells to measure apoptosis induction (**e**) Beads functionalised with each of the indicated SNAP-PhoCl-SpyC_1-6_ valency engineering covalent capture modules were subsequently functionalised with scFv-SpyTag, washed, and exposed to 405 nm light for 10 minutes. Samples were centrifuged, and 9 *μ*L of the supernatant loaded on a denaturing SDS-PAGE gel (**f**) Released multivalent assemblies from (**e**) were incubated with HeLa cells for 2 hours at the indicated concentrations. Cells were then assayed for apoptosis induction by incubation with NucView 488 and subsequent flow cytometry. Effective scFv concentration is the concentration of scFv in each well regardless of its multivalent state; data were obtained in triplicate; the dashed line indicates 50 % apoptosis; and the sigmoid curves are fitted Hill equations.

Many protein:protein interactions rely upon specific valencies of the interacting partners to trigger a specific cellular response^53,54^. Engineering the valency state of protein-based therapeutics that are designed to drug such biological systems typically relies upon laborious in-frame cloning and expression, limiting the capacity of a researcher to investigate many different drugs at many different valencies. The SpyCatcher technology has already been demonstrated to facilitate valency engineering through the post-translational assembly of monomeric nanobody-SpyTag into multivalent constructs via capture on SpyCatcher-coiled coil domain fusions^22^. We build upon this work by capturing SpyTag fusion proteins on PHD beads functionalised for valency engineering, thus taking advantage of surface immobilisation for washing and handling, and the subsequent release of assay components (e.g. in response to a supplied cue of light) to remove surface effects completely. To this end, we mounted distinct populations of beads with one of six SNAP-PhoCl-SpyCatcher fusion proteins (SNAP-PhoCl-SpyCatcher_1-6_ differing in the number of SpyCatcher repeats). Subsequent incubation with a monomeric SpyTag fusion protein results in assembly into photoreleasable, tunably multivalent constructs, depending only on the SpyCatcher module used (***Figure 6a***). An anti-TRAIL-R1 scFv^55^ (3B04) was chosen as a candidate for molecular engineering as related scFv TRAIL-R1 agonists^56^ reformatted as IgG had undergone clinical trial, with no clinical benefit seen in either non-small-cell lung cancer^57^ or colorectal cancer^58^. TRAIL-R1 is widely considered to signal as a trimer and, *in vivo*, is agonised by the trimeric TRAIL^59^, and we therefore hypothesised that enhanced potency could be achieved by engineering multivalent versions of the scFv. Similar multivalency engineering approaches have been carried out for nanobodies that target TRAIL-R2, a highly related receptor also found to be overexpressed on cancer cells, with great success^22,60^, but to our knowledge no such investigation has been carried out for scFvs targeting TRAIL-R1.

Initially we investigated the effect of making 3B04 trivalent (***Figure 6d***), through the incubation of 3B04-SpyTag with beads functionalised with SNAP-PhoCl-SpyCatcher_3_ and the subsequent exposure of half of these beads to 405 nm light. We observed that release of the multivalent assembly from the bead surface is necessary to fully induce apoptosis, presumably due to the sequestration of trivalent scFv assemblies within the volume of the bead, inaccessible to the cell surface receptors. It was straightforward to further engineer the valency state of 3B04-SpyT through incubation with separate bead populations, each functionalised with one of the six valency engineering modules (SNAP-PhoCl-SpyCatcher_1-6_). Subsequent exposure to 405 nm light released each of the fully 3B04-conjugated valency engineering modules into solution with high purity (***Figure 6e***). We incubated serial dilutions of each of these constructs (normalised to the effective scFv concentration) with HeLa cells for 2 hours and measured apoptosis induction using a fluorogenic caspase-3 substrate (NucView 488; ***Figure 6f***). Decreases in the EC50 values indicate significant increases in potency for all multivalent constructs over monovalent scFv, (e.g. >50-fold for the pentavalent versus monovalent format; ***Table 1***). Intriguingly, we observe an approximately two-fold *reduction* in potency when increasing scFv valency from 4x or 5x to 6x. This observation is consistent with previous observations that TRAIL-R1 signalling is dependent not only on trimerisation, but also on co-localisation of numerous TRAIL-R1 trimers within lipid rafts^61^. We speculate that the 4x and 5x constructs may promote trimer formation whilst also forming a lateral ‘bridge’ between consecutive TRAIL-R1 trimers, whereas the 6x construct may only enhance formation of a pair of trimers.

**Table 1.** EC50 values and standard deviations for multivalent antibody-induced cancer cell apoptosis. All EC50 values differ significantly from each other (p < 0.005, Welch’s two-tailed t-test). Conditions as per Figure 6f.

| scFv valency | EC50 (nM) |
| --- | --- |
| 1 | 114 ± 18 |
| 2 | 15.8 ± 1.2 |
| 3 | 8.21 ± 0.26 |
| 4 | 2.39 ± 0.17 |
| 5 | 1.99 ± 0.17 |
| 6 | 5.88 ± 0.52 |

## Conclusions and Implications

### An accessible, personalised technology platform for protein immobilisation

In contrast to commercial microbeads (made of e.g. polystyrene), PHD beads have user-definable attachment points and therefore bring customisable orthogonality and control over the valency of protein immobilisation into the hands of the researcher, who can exert this control ‘at home’ – in their (investigator’s) laboratory – simply by modifying the concentration of components in the hydrogel synthesis mixture. This: reduces reliance on commercial suppliers; avoids batch-to-batch variation outside the control of the researcher; enables a simple method for delivering user-defined amounts of protein to bioassays; and allows personalised variation of the type of tags used. Furthermore, the simple microfluidic bead synthesis ensures monodispersity at a level of control that is not available for commercial beads, providing flexibility and robustness to bioassays. Attachment points are selective (allowing e.g. direct purification of the protein from a cell lysate), which is brought about by covalent tagging – in addition to SNAP- and Halo-tag as used in this study, other tags are available^62^. The site-specific nature of protein capture minimises the potential impact of immobilisation on the activity of the protein of interest, whilst the covalent nature ensures that captured proteins remain stably associated with the hydrogel and do not leach into solution. Surface effects that are frequently encountered when proteins are physically immobilised on plastic surfaces are minimized, and hydrogels can be expected to mimic the natural environment for soluble proteins much better than a hydrophobic surface. The 3D distribution of attachment points throughout the hydrogel volume (rather than the surface of commercial microbeads), enables each bead to be decorated with 150 million protein molecules or more (∼1.5 billion for 500 *μ*M BG beads) in contrast with ∼660 thousand protein molecules captured on commercial streptavidin beads, ***Figure SI3b***. Finally, hydrogel beads are optically transparent, so that fluorescent measurements are possible and strong signal over background can be detected in all fluorescent channels, while commercial magnetic polystyrene beads exhibit autofluorescence in relevant channels, limiting assay sensitivity^16^.

### Versatile Assay Formatting

Based on the modular design principles of synthetic biology, PHD beads can be decorated by attaching tagged protein constructs in a generic way, in an effectively “plug and play” solution for biological experiments and engineering. This approach mirrors ‘click chemistry’^18^ by providing universal procedures for attachment that do not have to be adjusted on a case-by-case basis. Direct capture of POIs as SNAP or Halo-tag constructs initially simplifies protein purification directly from cell lysates, and this direct capture can be further augmented by the use of secondary capture modules which enable the expansion of protein capture to endogenous untagged targets (e.g. IgG) through the use of defined recombinant affinity reagents. We have developed a suite of these, focussing on bacterially expressible scaffolds to increase accessibility to the technology, and this suite could be readily expanded through the fusion of other affinity reagents (e.g. DARPins, nanobodies) to SNAP- or Halo-tag via modular cloning strategies. The use of defined, recombinant affinity reagents at the core of the PHD technology satisfies an urgent need to reduce the use of animal-derived, polyclonal reagents (as highlighted in recent EU directives^63^). Including secondary covalent capture modules (e.g. SpyCatcher/SpyTag, SnpCatcher/SnpTag) adds an extra layer of stable engineerability to the system and enables a second dimension of orthogonality for the creation of multifunctional hydrogels, while the use of valency-engineering modules allows monomeric proteins to be readily assembled into multivalent constructs. Complex multivalent and/or multi-protein decorations are accessible from (separate or mixed) solutions of monomers – these decorations are assembled on-bead and render cloning of additional multivalent constructs unnecessary. Multivalency^64,65^ and induced proximity^66^ is a natural mechanism of enhancing and manipulating interactions in biological systems by cooperativity.^67^ There are no general rules for the design of multivalent constructs that take advantage of entropic, avidity or co-localisation effects, so the orientation of monomers has to be empirically explored and an experimental format to empirically assess the contribution of multivalency is necessary. This fact is highlighted in our work by the most potent induction of apoptosis being a pentavalent antibody format despite knowledge that the target (TRAIL-R1) is agonised by a trimeric ligand *in vivo*. Typically, multivalent constructs are cloned and expressed as in-frame fusion proteins, requiring extensive and often practically difficult cloning (e.g. for sequence-homologous repeats that create PCR problems), alongside often expensive and complex mammalian cell expression (e.g. in the case of IgG), limiting both the accessibility of protein engineering and its throughput. However, with PHD beads judicious choice of valency engineering modules can bring about such constructs in multiple permutations simply by incubation instead of cloning, once the monomeric modules are available.

Versatility is further boosted by the possibility of photorelease. Steric hindrance and proximity to an ill-defined or hydrophobic surface can limit the applicability of protein assays on beads (in particular for cell-protein interactions), even though the 3D distribution in PHD beads and the solution-like nature of the hydrogel minimise these effects. However, the feature of controlled release of the bead-displayed proteins by optical control removes this common objection against the use of immobilised proteins in assays (as seen by the release of small molecule compounds in OBOC assays^68^). We show that trivalent scFv has to be released from beads in order to potently induce apoptosis. Future applications to take advantage of optical release will include e.g. functional tests with proteins that need to be internalised to target intracellular processes or the control of growth factor presentation for tissue engineering. Further controlled release mechanisms, such as protease sites could be added to modules, enabling for instance the tissue-specific release of sequestered/inactive protein drugs.^69,70^ Taken together, the versatility of PHD beads allows an unprecedented degree of freedom in the design of bioassay experiments; straightforward bead-mediated harvesting of proteins from lysates, valency control (both at the hydrogel decoration stage and for protein constructs), orthogonality of the coupling chemistry (through various tags) and controlled release constitute a technology suite capable of simplifying the planning and execution of discovery campaigns based on modularity (**Figure 7**). We have demonstrated the simple reformatting of beads and proteins for investigating protein:protein interactions, enzymatic catalysis, bacteriolysis and phenotypic assays, but an even wider range of assays and applications is conceivable and take advantage of salient features of PHD beads: biocatalysis, *in vivo* drug delivery, controlled release, and sensors.

**Figure 7.**
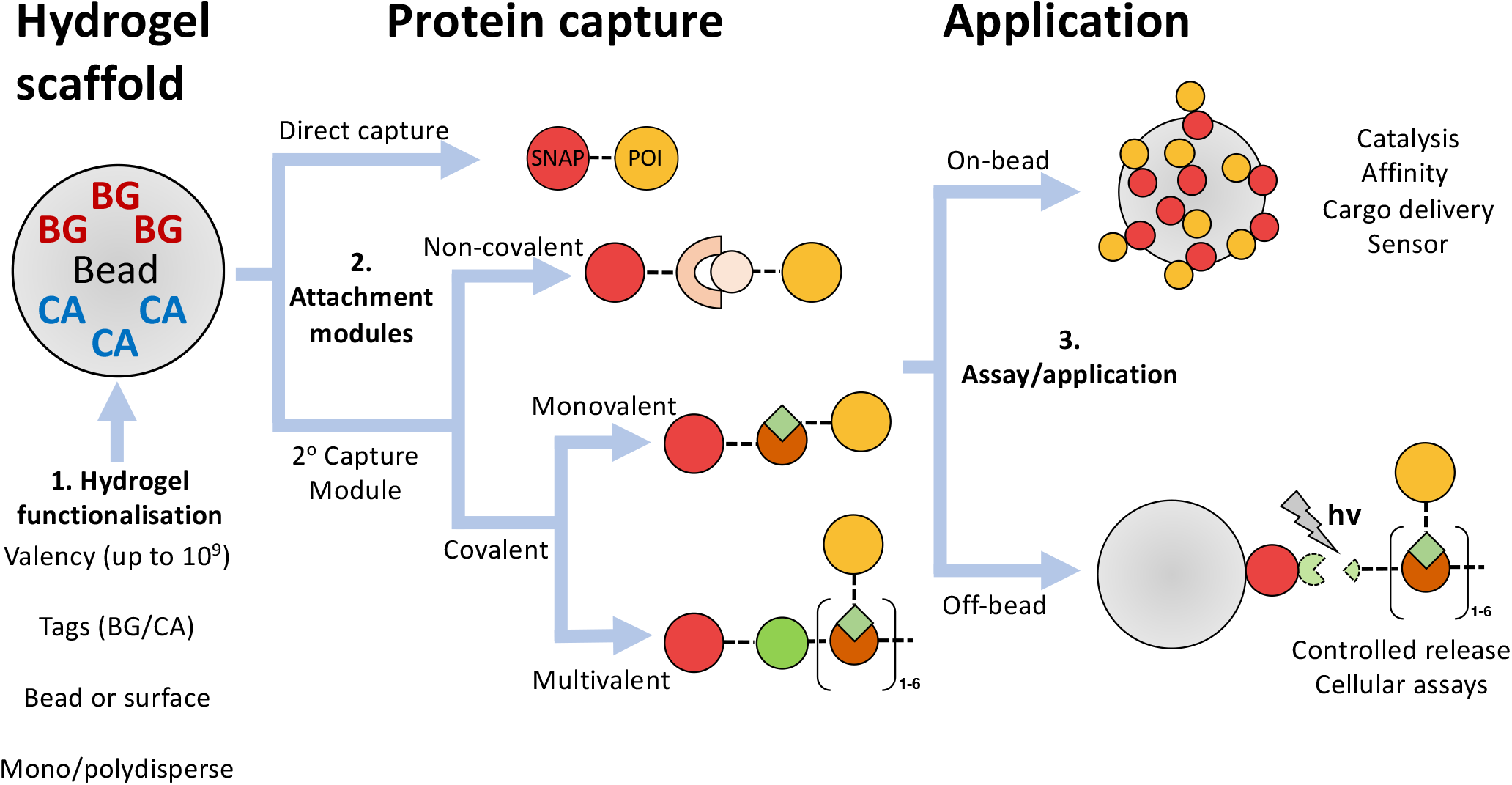
Overview of a modular “build-an-assay” strategy based on PHD beads. Starting from functionalised microbeads (**1**, *see below*), choices that define the assay format include the desired valency of each single bead as well as the loading of orthogonal protein capture into the system (controlled by the input concentrations of BG and CA). Next, one can choose how to capture a desired protein (**2**): either directly as a SNAP or Halo tag fusion protein, or via secondary capture modules. Secondary capture modules add the capability to specifically capture native or tagged proteins non-covalently, or to specifically and covalently capture proteins bearing tags, e.g. using the SpyTag-SpyCatcher or SnpTag/SnpCatcher technologies. At this stage one can also choose to create multivalent constructs from monomeric input proteins of interest through the use of valency engineering modules. Finally (**3**), the captured proteins can be tested in on-bead assays (e.g. for their affinity), or released from bead in response to irradiation of light, so that the new molecular assemblies can be assayed in solution (e.g. for phenotypic cellular assays). Monodisperse beads can be created in microfluidic devices via water-in-oil emulsions. The design of the microfluidic device and its operation determines the bead size. Alternatively, polydisperse emulsions protocols can be used to make beads at the price of a larger size distribution. As an alternative to the bead format, functionalised hydrogels can also be created on a surface (e.g. for cell culture).

## EXPERIMENTAL SECTION

### Protocol for hydrogel bead synthesis and functionalisation

1. The small molecule anchors (methacrylate-PEG-benzylguanine/-chloroalkane; Table S2.1) for hydrogel functionalisation were synthesised by mixing one volume of 40 mM BG-PEG-NH_2_ (NEB S9150S) *or* 40 mM chloroalkane-PEG-NH_2_ (Promega P6741) with one volume of 40 mM methacrylate-NHS (Sigma 730300) overnight at room temperature at 400 rpm in the presence of a 1.5-fold molar excess of triethylamine (Sigma 471283). All solutions were prepared fresh from powder in anhydrous DMSO (Merck 276855) except triethylamine which was added from neat stock. After overnight incubation the reaction was quenched with 3 volumes of 100 mM Tris-HCl (pH 8.0) and rolled 1 hour at room temperature, yielding a final concentration of 5 mM product.
2. To prepare functionalised beads, unpolymerised hydrogel mix (10 mM TrisHCl (pH 7.6), 1 mM EDTA, 15 mM NaCl, 6.2% (v/v) acrylamide, 0.18% (v/v) bisacrylamide, 0.3% (w/v) ammonium persulfate) containing the small molecule anchors was encapsulated in oil (008-Fluorosurfactant 1.35% w/w, RAN Biotechnologies, TEMED 0.4 % v/v in HFE-7500 (3M Novec)) in a microfluidic droplet generator (Figure S2.1), as previously described.^27^ After encapsulation the emulsion was incubated overnight at 65 °C under mineral oil. The next day, polymerised hydrogel beads were recovered by breaking the emulsion with 800 *μ*L wash buffer (100 mM Tris-HCl, 0.1 % Tween-20), and 200 *μ*L 1H,1H,2H,2H-perfluorooctanol (PFO, 97%, Alfa Aesar). The tube was inverted several times and briefly centrifuged for 5 seconds at 100 *g* before recovering the aqueous bead-containing phase into a fresh tube. Large polyacrylamide particles were removed by passing the mixture through a 10 *μ*m filter (CellTrics) for 30 seconds at 200 *g* before using a haemocytometer (KOVA Glasstic) to determine the ‘concentration’ of beads in the suspension. These beads are stable at 4 °C for at least one year. For all assays beads are typically incubated and washed in buffer (100 mM Tris-HCl, 0.1 % Tween-20). In other buffers and in unbuffered water, the bead pellet after centrifugation can be difficult to identify.
3. SNAP-tag/Halo-tag fusion proteins were captured by incubating with a defined number of beads for >30 minutes with rolling in wash buffer. After protein capture, beads were typically washed 3 times in wash buffer. Subsequent capture of tagged or untagged proteins was performed in the same manner.
4. On-bead photocleavage was carried out by attaching PCR tubes containing beads to a cooled metal block and exposing to 405 nm light at full power from a LED (M405L2 Thorlabs) driven by LEDD1b (Thorlabs).

## ASSOCIATED CONTENT

The Supporting Information is available free of charge at https://pubs.acs.org/doi/ and contains experimental results (HPLC analysis of BG formation, functionalisation of polydisperse PHD beads, valency calculations, cell lysate saturation binding and modular Gibson assembly design) and experimental procedures (benzylguanine analysis via HPLC), microfluidics and bead handling, protein expression and purification, molecular biology of module construction, mammalian cell culture, biological assays)

## AUTHOR INFORMATION

## AUTHOR CONTRIBUTIONS

T.F., J.D.R. and F.H. designed research, T.F., J.D.R. and C.M. performed research, T.F., J.D.R. and F.H. analyzed data, R.M. and F.H. directed research and T.F., J.D.R. and F.H. wrote the manuscript with feedback from all authors. All authors have given approval to the final version of the manuscript.

## FUNDING SOURCES

BBSRC iCASE studentship (to T. F., BB/M016692/1) and BBSRC-CTP studentship (J.D.R, BB/R505055/1). FH is an ERC Advanced Investigator (695669).

## ACKNOWLEDGMENT

The authors thank members of the Hollfelder group for comments on the manuscript, and J.D.F. Schnettler-Fernandez for the P91 vector and substrate. Thanks also to Dr. Joana Cerveira of the Cambridge BioPath Flow cytometry facility, Camilla Trevor (AstraZeneca), Josie Holstein (University of Cambridge) and the Core Tissue culture facility (AstraZeneca) for their guidance and practical support.

## Notes

### Competing Interest Statement

The authors have declared no competing interest.

