## Supplementary for "Gigavalent display of proteins on monodisperse polyacrylamide hydrogels as a versatile modular platform for functional assays and protein engineering"

**S1 Supplementary experimental figures and tables	S2**

Figure S1.1. Analysis of methacrylate-PEG-benzylguanine synthesis reaction products by UV HPLC

Table S1.1. Quantification of BG-PEG-MA (MA-BG) yield by substrate depletion

Figure S1.2. Creation of functionalised polydisperse PHD beads

Figure S1.3. Calculation of number of valencies per PHD bead

Figure S1.4. Capture of protein directly from cell lysate

Figure S1.5. Gibson assembly and restriction-ligation cloning strategies for module construction

**S2 Supplementary Materials and Methods	S6**

2.1 Analysis of benzylguanine-containing compounds (HPLC)

2.2 Microfluidic chip design, operation and bead handling

2.3 Protein expression and purification

2.4 Molecular biology of module construction

2.5 Mammalian cell culture

2.6 Fluorescent, enzymatic and phenotypic assays

**S3 References	S18**

**S1 Supplementary experimental figures and tables**

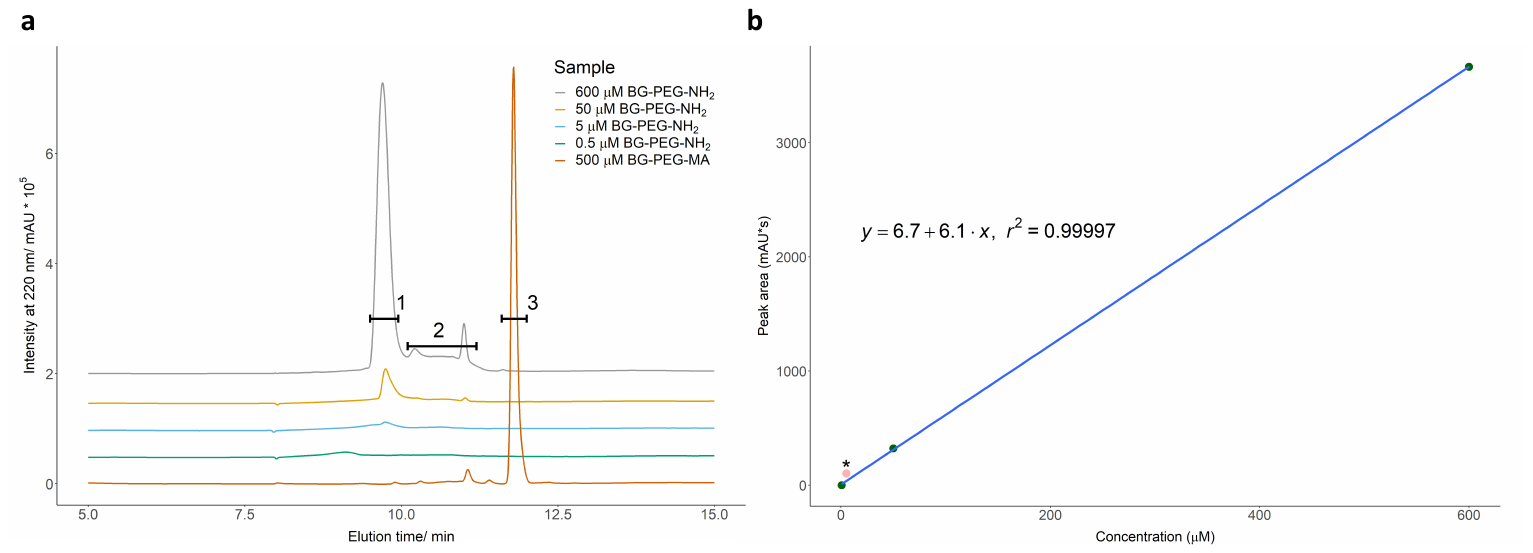

**Figure S1.1. Analysis of methacrylate-PEG-benzylguanine synthesis reaction products by UV HPLC. (a)** HPLC UV chromatograms of BG-PEG-NH_2_ standards and the reaction products of an overnight click reaction of BG-PEG-NH_2_ with MA-NHS. Peak identities were determined by comparison to control chromatograms of potential molecules (data not shown). Peak **1** was identified as BG-PEG-NH_2_ by its presence as the dominant molecule in commercially available BG-PEG-NH_2_ and its depletion upon addition of (reaction with) MA-NHS. Signals in region **2** most likely represents impurities or incomplete products of commercial BG-PEG-NH_2_ synthesis. Peak **3** appears only when reacting BG-PEG-NH_2_ with MA-NHS, and not in the HPLC chromatograms of either substrate individually, strongly suggesting it is MA-BG. In the absence of a pure standard of MA-BG, we estimated reaction efficiency by depletion of the BG-PEG-NH_2_ peak upon reaction with an equimolar amount of MA-NHS. Chromatogram baselines are offset from each other by 50000 mAU for visualisation purposes; there is no time offset. **(b)** BG-PEG-NH_2_ standard curve. Due to peak broadening and coalescence with those of the commercial impurities at low concentrations, we considered the peak integration of 5 µM BG-PEG-NH_2_ (*; pale red datapoint) to be anomalously high, so a standard line was fit through the peak areas of the 600 µM, 50 µM and 0.5 uM (peak area zero) BG-PEG-NH_2_ for estimation purposes. We note that excluding the omitted datapoint results in a lower estimate of reaction efficiency than including it. The fitted line and its equation are overlaid on the raw datapoints along with the r-squared value.

| Sample | Concentration (μM) | 9.6 - 10.0 min peak area (mAU*min) | BG-PEG-NH_2_ concentration estimate (μM) |
| --- | --- | --- | --- |
| BG-PEG-NH_2_ | 600 | 3665 |  |
| BG-PEG-NH_2_ | 50 | 322 |  |
| BG-PEG-NH_2_ | 5 | 103* |  |
| BG-PEG-NH_2_ | 0.5 | N/A |  |
| BG-PEG-NH_2_ | 500 | 10 | 0.57 |

**Table S1.1. Quantification of BG-PEG-MA (MA-BG) yield by substrate depletion.** Peak areas obtained by integration, and estimate of remaining BG-PEG-NH_2_ using the standard curve derived in ***Figure S1a***. The practical limitations outlined above mean that we can state with confidence that the reaction turnover is greater than 90 % product, but consider it very likely that the reaction is close to 99 % efficient based on our quantification – this estimate is also in very close agreement with our subsequent estimates of bead anchor density using SNAP-GFP titration (***Figure 2c*** and ***Figure S1.3***).

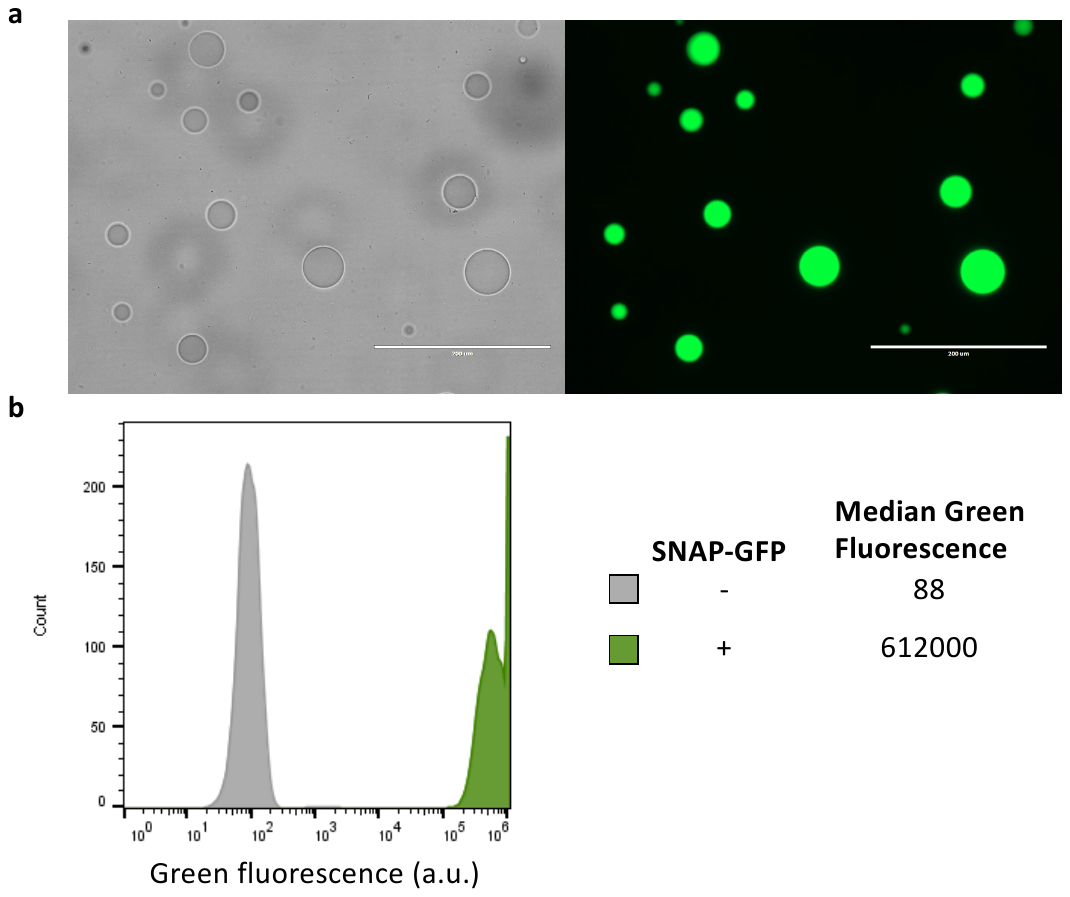

**Figure S1.2.** Creation of functionalised polydisperse PHD beads. Monomeric hydrogel mix (50 µM BG) and oil were setup as per standard protocol for flow-focusing (Main text, Methods) and, instead of encapsulation in a microfluidic chip, were vortexed at full power for 30 seconds. The resulting emulsion was then handled identically to the flow focussing protocol (**a**) Polydisperse beads were incubated with SNAP-GFP (10 µM, 100 µl), washed and analysed by fluorescent microscopy (left panel; brightfield, right panel: GFP channel). Scale bar shown corresponds to 200 µm. (**b**) Polydisperse beads were incubated +/- SNAP-GFP and analysed by flow cytometry. The same gain settings were used as for **Figure 2c**.

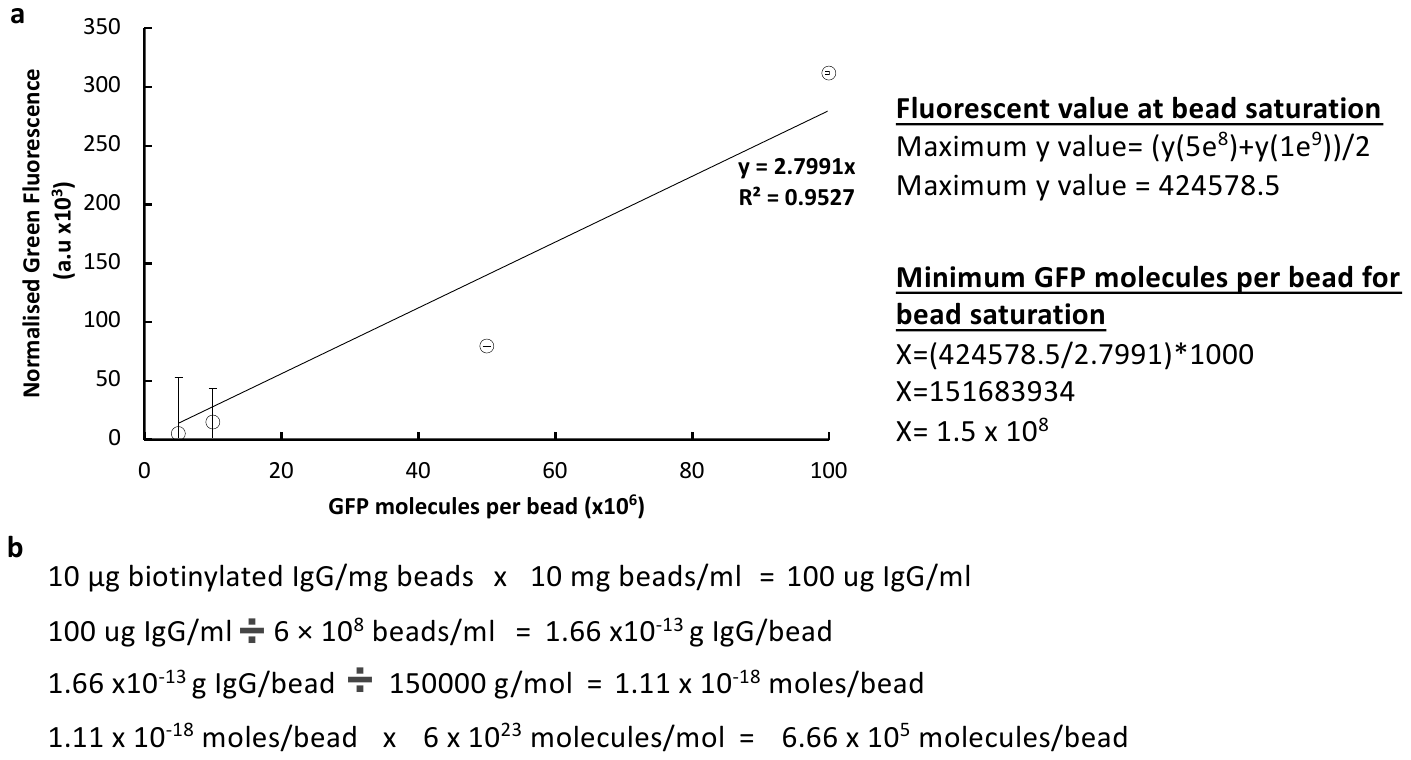

**Figure S1.3.** Calculation of number of valencies per PHD bead. (**a**) The linear section (5e^6^-1e^8^ GFP molecules per bead) of fluorescent data from **Figure 2c** was taken and a line of best fit calculated. This equation (y=2.7991x) was used to estimate the number of SNAP-GFP molecules required to saturate a bead based on the average maximum signal (for the y values of 5x10^8^ and 1x10^9^ SNAP-GFP molecules per bead). (**b**) Calculation of the valency of protein on the surface of M-280 Streptavidin Dynabeads suggests densities of <10^6^ molecules per bead (consistent with previous measurements, ref. 4).

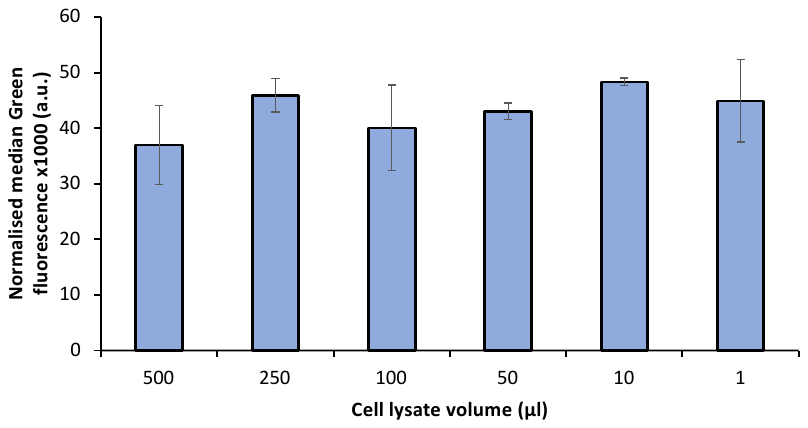

**Figure S1.4.** Capture of protein directly from cell lysate. *E. coli* cultures expressing SNAP-SnpT (**Table S2.2**) were grown in triplicate overnight in deepwell plates (1 ml per well). Cultures were pelleted and resuspended in 50 µL BugBuster (1x) for cell lysis. Bead wash buffer (450 µL) was added to each well after 1 hour of lysis, and the indicated volumes taken for incubation with fifty thousand 50 µM BG, Ø20 µm beads for 30 minutes, followed by washing (3 x 200 µl bead wash buffer) and incubation with GFP-SnpC (10 µM, 100µl). After washing, the beads were analysed by flow cytometry. The fluorescence values are the mean of triplicates after subtraction of the signal seen for 0 µM BG PHD beads.

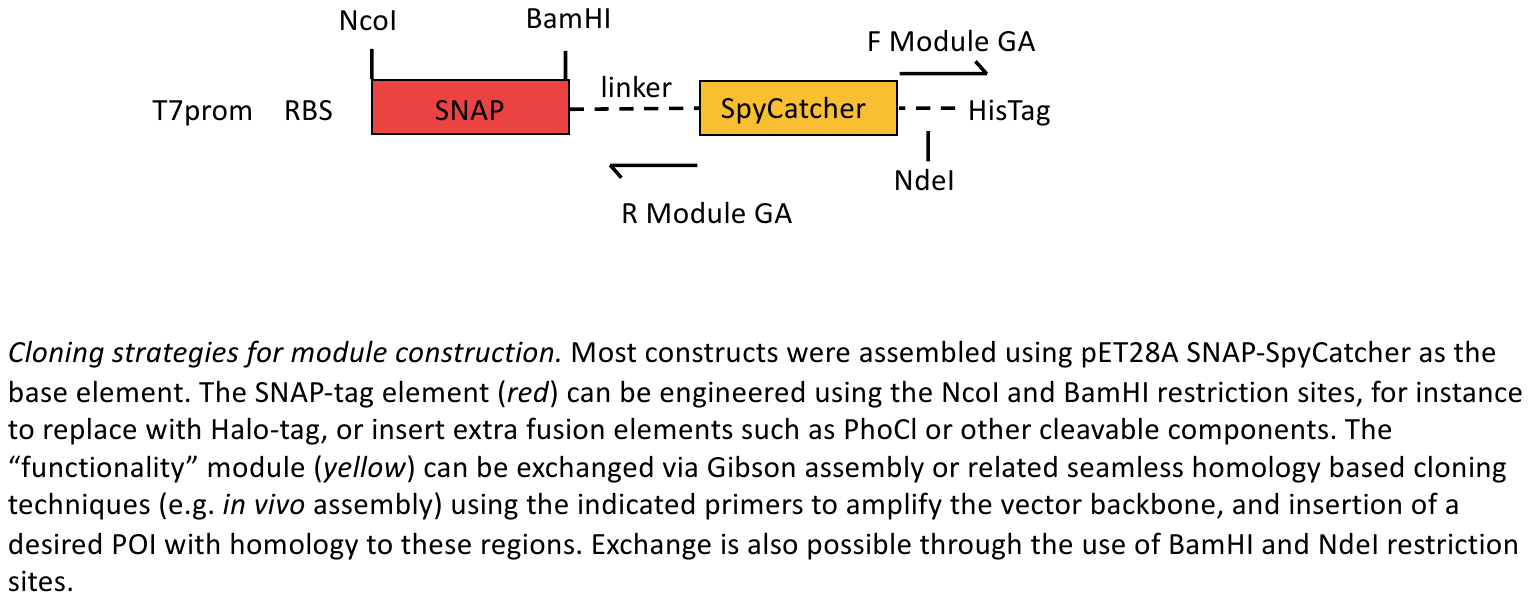

**Figure S1.5.** Gibson assembly and restriction-ligation cloning strategies for module construction. Modular cloning strategies are exemplified using pET28A SNAP-SpyCatcher (**Table S2.2**) as the base element. The SNAP-tag element (red) can be engineered using the NcoI and BamHI restriction sites, for instance to be replaced with Halo-tag, GFP, or extra fusion elements such as PhoCl or other cleavable components. The “functionality” module (yellow) can be exchanged via Gibson assembly or related seamless homology based cloning techniques (e.g. *in vivo* assembly) using the indicated primers (F and R Module GA) to amplify the vector backbone, and insertion of a desired POI with homology to these regions. Exchange is also possible through the use of BamHI and NdeI restriction sites.

**S2 Supplementary Materials and Methods**

**2.1 Analysis of benzylguanine-containing compounds (HPLC)**

BG-PEG-NH_2_ was obtained from NEB (S9150S) in powder form, and MA-NHS from Sigma (730300). Both vials were allowed to reach room temperature before resuspending all of the BG-PEG-NH_2_ to 40 mM in anhydrous DMSO (Merck 276855). An excess of 40 mM MA-NHS solution was prepared by weighing the powder and adding an appropriate volume of DMSO. Triethylamine was added to the BG-PEG-NH_2_ solution at a 1.5X molar amount before immediately adding a 1X molar amount of the MA-NHS solution, and the reaction mixture was shaken at 400 rpm, 30 °C overnight. A small aliquot of this mixture was taken, while the remainder was quenched with 3 volumes of 100 mM Tris-HCl (pH 8.0), rolling 1 hour at room temperature. The aliquot was diluted in MilliQ water to 500 µM MA-BG (assuming complete reaction) before analysis on a reversed phase C18 HPLC column (Fisher Scientific 12498357) using the following program: 5 min post-injection equilibration in water; gradient to 60 % acetonitrile over 10 min, then held for 3 min; gradient to 100 % acetonitrile over 2 min, held for 5 min; gradient to water over 5 min. The BG-PEG-NH_2_ standard curve was prepared fresh from powder and analysed in the same way.

**2.2 Microfluidic chip design, operation and bead handling**

The 20 µm flow-focussing microfluidic device (Figure S2.1; The CAD files for this, and other devices can be downloaded from our repository DropBase (<https://openwetware.org/wiki/DropBase:droplet_generation_2_inlets>)) was fabricated by standard soft lithography procedures using a high-resolution acetate mask (Microlithography Services Ltd.) and SU-8–2025 photoresist patterning. PDMS monomer and curing agent were mixed (10:1), degassed and then poured onto the chip master design prior to further degassing. After PDMS solidification (65 °C, O/N), the chip was cut out and inlets and outlets punched through. Subsequently the chip was exposed to an oxygen plasma and sealed onto a microscope glass slide. Hydrophobic modification of the channel surfaces was achieved by injecting a solution of 1% (v/v) trichloro(1H,1H,2H,2H-perfluorooctyl)silane (Sigma) in HFE-7500 oil into the channels.

Ø20 µm hydrogel beads are made at a rate of ~8 kHz by flowing an aqueous stream of unpolymerized hydrogel mix at 2 µL/min and sheath fluid at 5 µL/min through the 20 µm flow-focussing chip (Table S2.1). Gas-tight glass syringes (SGE) were driven by syringe pumps (Nemesys) and connected to the chip via fine pore PTFE tubing (ID 0.38 mm, OD 1.09 mm, Smith Medical). After encapsulation the emulsion is incubated overnight at 65 °C, overlaid by 200 µl mineral oil. The next day, polymerised hydrogel beads are recovered by removing the mineral oil by pipetting and breaking the emulsion with 800 µL wash buffer (100 mM Tris-HCl, 0.1 % Tween-20) and 200 µL 1H,1H,2H,2H-perfluorooctanol (PFO, 97%, Alfa Aesar). The tube is inverted several times until the emulsion is broken, and briefly centrifuged for 5 seconds at 100 *g*. The aqueous bead-containing phase is recovered into a fresh tube. Large polyacrylamide particles are removed by passing the mixture through a 10 µm filter (CellTrics) for 30 seconds at 200 *g*. Hydrogel beads are then counted on a haemocytometer (KOVA Glasstic). These beads can be stored at 4 °C. For all assays, beads were typically incubated and washed in wash buffer.

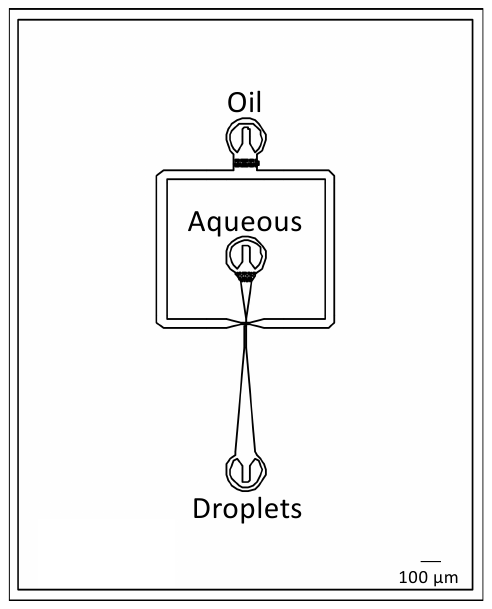

**Figure S2.1** Microfluidic device for 20 µm PHD bead synthesis. Aqueous hydrogel mix is flowed at 2 µl/min, and oil at 5 µl/min.

**Table S2.1.** Hydrogel mix, sheath fluid and buffer composition for production and handling of functionalised polyacrylamide hydrogel beads.

| Mixture | Material and stock concentration | Volume (µL) |
| --- | --- | --- |
| Unpolymerised hydrogel mix | Tris-HCl (100 mM, pH 8.0)  NaCl (5 M)  EDTA (0.5 M)  Acrylamide (40 % v/v)  Bis-acrylamide (2 % v/v)  APS (10 % w/v)  Acrydite-benzyl guanine (5 mM)  Acrydite-chloroalkane (5 mM)  Water | 50  1.5  1  75  47.5  15  Variable  Variable  To 500 µL |
| Sheath fluid for encapsulation | 008- Fluorosurfactant (10 % w/w RAN Biotechnologies)  TEMED (neat)  HFE-7500 (3M Novec) | 135  4  861 |

**2.3 Protein expression and purification**

**2.3.1 *E. coli* expression of fusion proteins**

All proteins (except scFv-SpyTag and P91-SpyTag) were expressed in BL21 (DE3) cells. Briefly, 5 mL overnight cultures were inoculated into 750 mL 2YT media with appropriate selection antibiotics in baffled flasks, and grown at 37 °C for 2.5 hours at 200 rpm, then IPTG was added to a final concentration of 500 µM and cells grown overnight at 25 °C.

For protein extractions cells were pelleted, washed in 25 mL PBS and resuspended in 10 mL of Lysis buffer (1 x BugBuster, 0.2 cOmplete EDTA free protease inhibitor tablets, 1 µL Benzonase) and incubated with rolling at room temperature for at least 1 hour. Cell debris was pelleted, and the supernatant used for purification on 5 mL Super Ni-NTA Agarose Resin (Generon, NB-45-00042-25). The columns were washed with protein wash buffer (20 ml, 500 mM NaCl, 30 mM imidazole, 20 mM Tris-HCl, pH 8.0) and eluted with elution buffer (5 mL, 500 mM NaCl, 500 mM imidazole, 20 mM Tris-HCl, pH 8.0). Proteins were then concentrated and buffer exchanged into PBS using Amicon Ultra centrifuge filters with the appropriate molecular weight cutoffs. Protein concentration was then measured using A^280 nm^ values, and A^488 nm^ values (for GFP containing constructs) on a Nanodrop 2000 instrument. Proteins were stored at -80 °C long term, and 4 °C after thawing and for short-term usage.

**2.3.2 *In vitro* expression of P91-SpyTag**

P91-SpyTag was expressed from pIVEX P91-SpyTag using a standard 25 µl reaction setup of PURExpress (NEB E6800S) for 16 hours at 30^o^C. The reaction was then incubated with 812500 beads functionalised with SNAP-SpyCatcher for 1 hour with shaking at room temperature.

**2.3.3 Expression of scFv-SpyTag in CHO Cells**

CD33-3B04 scFv-SpyTag-HA-His (**Table S2.2**), codon-optimised for expression in CHO cells, was synthesised commercially by GeneArt and subcloned into pDest12.2 OriP using standard restriction cloning techniques with NotI-HF and NheI-HF (NEB). Vectors were amplified in *E. coli* (Zymo Research T3007) under ampicillin selection and extracted using a MidiPrep kit (Qiagen)

A 500 mL culture of high-yielding CHO cells^5^ was transfected with 11.75 mL of a solution containing 21 µg/µL vector DNA, 170 µg/mL PEI MAX (Polysciences Inc. 247651) and 150 mM NaCl. CHO cells were grown at 34 °C, 5 % CO_2_, ambient O_2_, shaking 150 rpm. Cultures were fed with a MedImmune proprietary nutrient supplement 4 hours after transfection and again after three days.

Seven days after transfection, cells were pelleted 30 min at 5000 *g*, 4 °C. The scFv was purified from the cell supernatant on an AKTA using a HisTrap Excel column (Cytiva 17371206) pre-equilibrated with 5 column volumes (CV) of PBS, washed post-injection with 3 CV 20 mM imidazole in PBS (adjusted to pH 7.4 with HCl), then eluted in 1 mL fractions using 5 CV 400 mM imidazole (pH 7.4, HCl). Flow rate 5 mL/min. Fractions were selected based on the UV absorbance chromatogram (A^280 nm^) peak, concentrated (Cytiva 28-9323-60) and buffer-exchanged (Cytiva 17085101) into PBS. Final protein concentrations were determined based on a Nanodrop A^280 nm^ reading and an extinction coefficient calculated using Benchling. Protein purity was analysed by Coomassie-stained SDS-PAGE, with only the expected band observed. The buffer-exchanged protein was stored at 4 °C until use and appears stable under these conditions for at least a year.

**Table S2.2.** DNA sequences for all vectors and inserts.

| Construct | Sequence |
| --- | --- |
| CD33-3B04 (scFv-SpyT-HA-His) | ATGCCTTTGCTTTTGCTGCTGCCTTTGTTGTGGGCTGGCGCTCTGGCTGAAGTGCAGTTGGTTCAGTCTGGCGCCGAAGTGAAGATGCCTGGCGCTTCCGTGAAGCTGTCCTGCAGAGTGTCTGGCGATACCTTCACCGCCTACTTCATCCACTGGGTCCGACAGGCTCCAGGACAGGGACTTGAATGGATGGGCTGGTTCAACCCCATCTCTGGCACCGCTGGATCCGCCGAGAAGTTCAGAGGTAGAGTGGCCATGACCAGAGACACCTCCATCTCTACCGCCTACATGGAACTGAACCGGCTGACCTTCGACGACACCGCCGTGTACTACTGTGCCAGACAGCACCGGGGCAACACCTTTGATCCTTGGGGACAGGGCACCCTGGTCACAGTTTCTAGCGGAGGCGGAGGATCTGGTGGTGGTGGATCTGGCGGCGGAGGTTCTGCTCAGTCTGCTCTGACACAGCCCGTGTCCGTGTCTGGATCTCCCGGCCAGTCTATCACCATCAGCTGTACCGGCACCTCCTCTGACATCGGCGCCTACAAATACGTGTCCTGGTATCAGCAGCACCCCGGCAAGGCTCCTAAGCTGGTCATCTACGAGGTGTCCAACCGGCCTTCTGGCGTGTCCTCTAGATTCTCCGGCTCCAAGTCCGGCCAGACCGCTTCTCTGACAATCAGCGGACTGCAGGCCGACGACGAGGCCGACTACTACTGCAATTCCTACCAGGGCTACAACACCTGGGTGTTCGGCGGAGGCACCAAAGTGACAGTTCTGGGAGCTGCTGCAGGCGCTTCTGGTGCTGGTGCTCATATCGTGATGGTGGACGCCTACAAGCCCACCAAAGGCTCTGGCTCTGGCAGATACCCCTACGACGTGCCCGATTACGCCAGCCACCACCATCATCACCACTGA |
| pIVEX P91-SpyTag | GAGGGCTTACCATCTGGCCCCAGTGCTGCAATGATACCGCGAGACCCACGCTCACCGGCTCCAGATTTATCAGCAATAAACCAGCCAGCCGGAAGGGCCGAGCGCAGAAGTGGTCCTGCAACTTTATCCGCCTCCATCCAGTCTATTAATTGTTGCCGGGAAGCTAGAGTAAGTAGTTCGCCAGTTAATAGTTTGCGCAACGTTGTTGCCATTGCTACAGGCATCGTGGTGTCACGCTCGTCGTTTGGTATGGCTTCATTCAGCTCCGGTTCCCAACGATCAAGGCGAGTTACATGATCCCCCATGTTGTGCAAAAAAGCGGTTAGCTCCTTCGGTCCTCCGATCGTTGTCAGAAGTAAGTTGGCCGCAGTGTTATCACTCATGGTTATGGCAGCACTGCATAATTCTCTTACTGTCATGCCATCCGTAAGATGCTTTTCTGTGACTGGTGAGTACTCAACCAAGTCATTCTGAGAATAGTGTATGCGGCGACCGAGTTGCTCTTGCCCGGCGTCAATACGGGATAATACCGCGCCACATAGCAGAACTTTAAAAGTGCTCATCATTGGAAAACGTTCTTCGGGGCGAAAACTCTCAAGGATCTTACCGCTGTTGAGATCCAGTTCGATGTAACCCACTCGTGCACCCAACTGATCTTCAGCATCTTTTACTTTCACCAGCGTTTCTGGGTGAGCAAAAACAGGAAGGCAAAATGCCGCAAAAAAGGGAATAAGGGCGACACGGAAATGTTGAATACTCATACTCTTCCTTTTTCAATATTATTGAAGCATTTATCAGGGTTATTGTCTCATGAGCGGATACATATTTGAATGTATTTAGAAAAATAAACAAATAGGGGTTCCGCGCACATTTCCCCGAAAAGTGCCACCTGACGTCTAAGAAACCATTATTATCATGACATTAACCTATAAAAATAGGCGTATCACGAGGCCCTTTCGTCTCGCGCGTTTCGGTGATGACGGTGAAAACCTCTGACACATGCAGCTCCCGGAGACGGTCACAGCTTGTCTGTAAGCGGATGCCGGGAGCAGACAAGCCCGTCAGGGCGCGTCAGCGGGTGTTGGCGGGTGTCGGGGCTGGCTTAACTATGCGGCATCAGAGCAGATTGTACTGAGAGTGCACCATATATGCGGTGTGAAATACCGCACAGATGCGTAAGGAGAAAATACCGCATCAGGCGCCATTCGCCATTCAGGCTGCGCAACTGTTGGGAAGGGCGATCGGTGCGGGCCTCTTCGCTATTACGCCAGCTGGCGAAAGGGGGATGTGCTGCAAGGCGATTAAGTTGGGTAACGCCAGGGTTTTCCCAGTCACGACGTTGTAAAACGACGGCCAGTGCCAAGCTTGCATGCAAGGAGATGGCGCCCAACAGTCCCCCGGCCACGGGGCCTGCCACCATACCCACGCCGAAACAAGCGCTCATGAGCCCGAAGTGGCGAGCCCGATCTTCCCCATCGGTGATGTCGGCGATATAGGCGCCAGCAACCGCACCTGTGGCGCCGGTGATGCCGGCCACGATGCGTCCGGCGTAGAGGATCGAGATCTCGATCCCGCGAAATTAATACGACTCACTATAGGGAGACCACAACGGTTTCCCTCTAGAAATAATTTTGTTTAACTTTAAGAAGGAGATATACATATGGCTAGCTGGAGCCACCCGCAGTTCGAAAAAGGCGCCATGACAGCAAGAAAAGTCGACTACACAGACGGTGCAACCCGCTGTATCGGTGAGTTTCATTGGGATGAAGGCAAGTCGGGCCCGCGTCCCGGCGTGGTGGTCTTTCCCGAGGCTTTCGGCCTCAACGACCATGCCAAGGAGCGCGCGCGGCGCCTTGCCGACCTCGGCTTTGCAGCCCTGGCGGCGGATATGCACGGAGACGCCCAGGTTTTCGATGCGGCGAGTCTCTCATCAACCATACAGGGCTACTACGGCGACCGCGCCCACTGGCGACGTCGTGCGCAGGCAGCGCTCGATGCACTGACGGCACAGCCAGAGGTGGACGGCAGCAAGGTGGCGGCCATCGGCTTTTGTTTCGGCGGTGCCACCTGCCTTGAACTGGCCCGCACAGGTGCGCCGCTGACCGCCATTGTCACCTTCCACGGCGGTTTGCTGCCGGAGATGGCAGGCGATGCCGGACGGATCCAGTCCAGTGTTCTGGTGTGCCATGGCGCTGATGATCCGCTCGTACAGGACGAAACCATGAAGGCCGTCATGGACGAGTTTCGTCGCGACAAGGTGGATTGGCAGGTGCTCTACCTCGGAAATGCGGTACACAGTTTCACCGATCCACTCGCTGGCAGTCACGGCATACCCGGGCTGGCCTATGACGCCACTGCCGAAGCCCGGTCGTGGACGGCCATGTGCAATCTGTTCAGTGAACTGTTCGGCAAAGGTGCGGCAGCAGGGGCCTCGGGGGCCGGAGCCCACATCGTGATGGTGGACGCCTACAAGCCGACGAAGGGTAGTGGAAGCGGCCGCTATCCGTATGATGTACCAGATTATGCAAGCCTCTAATAGGATCCGGCTGCTAACAAAGCCCGAAAGGAAGCTGAGTTGGCTGCTGCCACCGCTGAGCAATAACTAGCATAACCCCTTGGGGCCTCTAAACGGGTCTTGAGGGGTTTTTTGCTGAAAGGAGGAACTATATCCGGATATCCACAGGACGGGTGTGGTCGCCATGATCGCGTAGTCGATAGTGGCTCCAAGTAGCGAAGCGAGCAGGACTGGGCGGCGGCCAAAGCGGTCGGACAGTGCTCCGAGAACGGGTGCGCATAGAAATTGCATCAACGCATATAGCGCTAGCAGCACGCCATAGTGACTGGCGATGCTGTCGGAATGGACGATATCCCGCAAGAGGCCCGGCAGTACCGGCATAACCAAGCCTATGCCTACAGCATCCAGGGTGACGGTGCCGAGGATGACGATGAGCGCATTGTTAGATTTCATACACGGTGCCTGACTGCGTTAGCAATTTAACTGTGATAAACTACCGCATTAAAGCTTATCGATGATAAGCTGTCAAACATGAGAATTCGTAATCATGTCATAGCTGTTTCCTGTGTGAAATTGTTATCCGCTCACAATTCCACACAACATACGAGCCGGAAGCATAAAGTGTAAAGCCTGGGGTGCCTAATGAGTGAGCTAACTCACATTAATTGCGTTGCGCTCACTGCCCGCTTTCCAGTCGGGAAACCTGTCGTGCCAGCTGCATTAATGAATCGGCCAACGCGCGGGGAGAGGCGGTTTGCGTATTGGGCGCTCTTCCGCTTCCTCGCTCACTGACTCGCTGCGCTCGGTCGTTCGGCTGCGGCGAGCGGTATCAGCTCACTCAAAGGCGGTAATACGGTTATCCACAGAATCAGGGGATAACGCAGGAAAGAACATGTGAGCAAAAGGCCAGCAAAAGGCCAGGAACCGTAAAAAGGCCGCGTTGCTGGCGTTTTTCCATAGGCTCCGCCCCCCTGACGAGCATCACAAAAATCGACGCTCAAGTCAGAGGTGGCGAAACCCGACAGGACTATAAAGATACCAGGCGTTTCCCCCTGGAAGCTCCCTCGTGCGCTCTCCTGTTCCGACCCTGCCGCTTACCGGATACCTGTCCGCCTTTCTCCCTTCGGGAAGCGTGGCGCTTTCTCATAGCTCACGCTGTAGGTATCTCAGTTCGGTGTAGGTCGTTCGCTCCAAGCTGGGCTGTGTGCACGAACCCCCCGTTCAGCCCGACCGCTGCGCCTTATCCGGTAACTATCGTCTTGAGTCCAACCCGGTAAGACACGACTTATCGCCACTGGCAGCAGCCACTGGTAACAGGATTAGCAGAGCGAGGTATGTAGGCGGTGCTACAGAGTTCTTGAAGTGGTGGCCTAACTACGGCTACACTAGAAGGACAGTATTTGGTATCTGCGCTCTGCTGAAGCCAGTTACCTTCGGAAAAAGAGTTGGTAGCTCTTGATCCGGCAAACAAACCACCGCTGGTAGCGGTGGTTTTTTTGTTTGCAAGCAGCAGATTACGCGCAGAAAAAAAGGATCTCAAGAAGATCCTTTGATCTTTTCTACGGGGTCTGACGCTCAGTGGAACGAAAACTCACGTTAAGGGATTTTGGTCATGAGATTATCAAAAAGGATCTTCACCTAGATCCTTTTAAATTAAAAATGAAGTTTTAAATCAATCTAAAGTATATATGAGTAAACTTGGTCTGACAGTTACCAATGCTTAATCAGTGAGGCACCTATCTCAGCGATCTGTCTATTTCGTTCATCCATAGTTGCCTGACTCCCCGTCGTGTAGATAACTACGATACGG |
| pET SNAP-SpyCatcher | TCACCTGAATCAGGATATTCTTCTAATACCTGGAATGCTGTTTTCCCGGGGATCGCAGTGGTGAGTAACCATGCATCATCAGGAGTACGGATAAAATGCTTGATGGTCGGAAGAGGCATAAATTCCGTCAGCCAGTTTAGTCTGACCATCTCATCTGTAACATCATTGGCAACGCTACCTTTGCCATGTTTCAGAAACAACTCTGGCGCATCGGGCTTCCCATACAATCGATAGATTGTCGCACCTGATTGCCCGACATTATCGCGAGCCCATTTATACCCATATAAATCAGCATCCATGTTGGAATTTAATCGCGGCCTAGAGCAAGACGTTTCCCGTTGAATATGGCTCATAACACCCCTTGTATTACTGTTTATGTAAGCAGACAGTTTTATTGTTCATGACCAAAATCCCTTAACGTGAGTTTTCGTTCCACTGAGCGTCAGACCCCGTAGAAAAGATCAAAGGATCTTCTTGAGATCCTTTTTTTCTGCGCGTAATCTGCTGCTTGCAAACAAAAAAACCACCGCTACCAGCGGTGGTTTGTTTGCCGGATCAAGAGCTACCAACTCTTTTTCCGAAGGTAACTGGCTTCAGCAGAGCGCAGATACCAAATACTGTCCTTCTAGTGTAGCCGTAGTTAGGCCACCACTTCAAGAACTCTGTAGCACCGCCTACATACCTCGCTCTGCTAATCCTGTTACCAGTGGCTGCTGCCAGTGGCGATAAGTCGTGTCTTACCGGGTTGGACTCAAGACGATAGTTACCGGATAAGGCGCAGCGGTCGGGCTGAACGGGGGGTTCGTGCACACAGCCCAGCTTGGAGCGAACGACCTACACCGAACTGAGATACCTACAGCGTGAGCTATGAGAAAGCGCCACGCTTCCCGAAGGGAGAAAGGCGGACAGGTATCCGGTAAGCGGCAGGGTCGGAACAGGAGAGCGCACGAGGGAGCTTCCAGGGGGAAACGCCTGGTATCTTTATAGTCCTGTCGGGTTTCGCCACCTCTGACTTGAGCGTCGATTTTTGTGATGCTCGTCAGGGGGGCGGAGCCTATGGAAAAACGCCAGCAACGCGGCCTTTTTACGGTTCCTGGCCTTTTGCTGGCCTTTTGCTCACATGTTCTTTCCTGCGTTATCCCCTGATTCTGTGGATAACCGTATTACCGCCTTTGAGTGAGCTGATACCGCTCGCCGCAGCCGAACGACCGAGCGCAGCGAGTCAGTGAGCGAGGAAGCGGAAGAGCGCCTGATGCGGTATTTTCTCCTTACGCATCTGTGCGGTATTTCACACCGCATATATGGTGCACTCTCAGTACAATCTGCTCTGATGCCGCATAGTTAAGCCAGTATACACTCCGCTATCGCTACGTGACTGGGTCATGGCTGCGCCCCGACACCCGCCAACACCCGCTGACGCGCCCTGACGGGCTTGTCTGCTCCCGGCATCCGCTTACAGACAAGCTGTGACCGTCTCCGGGAGCTGCATGTGTCAGAGGTTTTCACCGTCATCACCGAAACGCGCGAGGCAGCTGCGGTAAAGCTCATCAGCGTGGTCGTGAAGCGATTCACAGATGTCTGCCTGTTCATCCGCGTCCAGCTCGTTGAGTTTCTCCAGAAGCGTTAATGTCTGGCTTCTGATAAAGCGGGCCATGTTAAGGGCGGTTTTTTCCTGTTTGGTCACTGATGCCTCCGTGTAAGGGGGATTTCTGTTCATGGGGGTAATGATACCGATGAAACGAGAGAGGATGCTCACGATACGGGTTACTGATGATGAACATGCCCGGTTACTGGAACGTTGTGAGGGTAAACAACTGGCGGTATGGATGCGGCGGGACCAGAGAAAAATCACTCAGGGTCAATGCCAGCGCTTCGTTAATACAGATGTAGGTGTTCCACAGGGTAGCCAGCAGCATCCTGCGATGCAGATCCGGAACATAATGGTGCAGGGCGCTGACTTCCGCGTTTCCAGACTTTACGAAACACGGAAACCGAAGACCATTCATGTTGTTGCTCAGGTCGCAGACGTTTTGCAGCAGCAGTCGCTTCACGTTCGCTCGCGTATCGGTGATTCATTCTGCTAACCAGTAAGGCAACCCCGCCAGCCTAGCCGGGTCCTCAACGACAGGAGCACGATCATGCGCACCCGTGGGGCCGCCATGCCGGCGATAATGGCCTGCTTCTCGCCGAAACGTTTGGTGGCGGGACCAGTGACGAAGGCTTGAGCGAGGGCGTGCAAGATTCCGAATACCGCAAGCGACAGGCCGATCATCGTCGCGCTCCAGCGAAAGCGGTCCTCGCCGAAAATGACCCAGAGCGCTGCCGGCACCTGTCCTACGAGTTGCATGATAAAGAAGACAGTCATAAGTGCGGCGACGATAGTCATGCCCCGCGCCCACCGGAAGGAGCTGACTGGGTTGAAGGCTCTCAAGGGCATCGGTCGAGATCCCGGTGCCTAATGAGTGAGCTAACTTACATTAATTGCGTTGCGCTCACTGCCCGCTTTCCAGTCGGGAAACCTGTCGTGCCAGCTGCATTAATGAATCGGCCAACGCGCGGGGAGAGGCGGTTTGCGTATTGGGCGCCAGGGTGGTTTTTCTTTTCACCAGTGAGACGGGCAACAGCTGATTGCCCTTCACCGCCTGGCCCTGAGAGAGTTGCAGCAAGCGGTCCACGCTGGTTTGCCCCAGCAGGCGAAAATCCTGTTTGATGGTGGTTAACGGCGGGATATAACATGAGCTGTCTTCGGTATCGTCGTATCCCACTACCGAGATATCCGCACCAACGCGCAGCCCGGACTCGGTAATGGCGCGCATTGCGCCCAGCGCCATCTGATCGTTGGCAACCAGCATCGCAGTGGGAACGATGCCCTCATTCAGCATTTGCATGGTTTGTTGAAAACCGGACATGGCACTCCAGTCGCCTTCCCGTTCCGCTATCGGCTGAATTTGATTGCGAGTGAGATATTTATGCCAGCCAGCCAGACGCAGACGCGCCGAGACAGAACTTAATGGGCCCGCTAACAGCGCGATTTGCTGGTGACCCAATGCGACCAGATGCTCCACGCCCAGTCGCGTACCGTCTTCATGGGAGAAAATAATACTGTTGATGGGTGTCTGGTCAGAGACATCAAGAAATAACGCCGGAACATTAGTGCAGGCAGCTTCCACAGCAATGGCATCCTGGTCATCCAGCGGATAGTTAATGATCAGCCCACTGACGCGTTGCGCGAGAAGATTGTGCACCGCCGCTTTACAGGCTTCGACGCCGCTTCGTTCTACCATCGACACCACCACGCTGGCACCCAGTTGATCGGCGCGAGATTTAATCGCCGCGACAATTTGCGACGGCGCGTGCAGGGCCAGACTGGAGGTGGCAACGCCAATCAGCAACGACTGTTTGCCCGCCAGTTGTTGTGCCACGCGGTTGGGAATGTAATTCAGCTCCGCCATCGCCGCTTCCACTTTTTCCCGCGTTTTCGCAGAAACGTGGCTGGCCTGGTTCACCACGCGGGAAACGGTCTGATAAGAGACACCGGCATACTCTGCGACATCGTATAACGTTACTGGTTTCACATTCACCACCCTGAATTGACTCTCTTCCGGGCGCTATCATGCCATACCGCGAAAGGTTTTGCGCCATTCGATGGTGTCCGGGATCTCGACGCTCTCCCTTATGCGACTCCTGCATTAGGAAGCAGCCCAGTAGTAGGTTGAGGCCGTTGAGCACCGCCGCCGCAAGGAATGGTGCATGCAAGGAGATGGCGCCCAACAGTCCCCCGGCCACGGGGCCTGCCACCATACCCACGCCGAAACAAGCGCTCATGAGCCCGAAGTGGCGAGCCCGATCTTCCCCATCGGTGATGTCGGCGATATAGGCGCCAGCAACCGCACCTGTGGCGCCGGTGATGCCGGCCACGATGCGTCCGGCGTAGAGGATCGAGATCTCGATCCCGCGAAATTAATACGACTCACTATAGGGGAATTGTGAGCGGATAACAATTCCCCTCTAGAAATAATTTTGTTTAACTTTAAGAAGGAGATATACCATGGACAAGGATTGTGAAATGAAACGCACCACACTGGACAGCCCTTTGGGGAAGCTGGAGCTGTCTGGTTGTGAGCAGGGTCTGCACCGTATAATTTTTCTGGGCAAGGGGACGTCTGCAGCTGATGCCGTGGAGGTCCCAGCCCCCGCTGCGGTTCTCGGAGGTCCGGAGCCCCTGATGCAGGCTACAGCCTGGCTGAATGCCTATTTCCACCAGCCCGAGGCTATCGAAGAGTTCCCCGTGCCGGCTCTTCACCATCCCGTTTTCCAGCAAGAGTCGTTCACCAGACAGGTGTTATGGAAGCTGCTGAAGGTTGTGAAATTCGGAGAAGTGATTTCTTACTCTCACTTAGCAGCCCTGGCAGGCAACCCCGCAGCCACGGCAGCAGTGAAGACGGCACTGAGTGGCAATCCTGTCCCTATCCTGATCCCGTGCCACAGAGTGGTCCAGGGGGATTTGGATGTGGGCGGTTACGAGGGTGGACTGGCCGTGAAGGAATGGCTTCTGGCCCATGAAGGCCACCGGTTGGGGAAGCCAGGCTTGGGATCCGGTTCCGAAACGCCTGGAACGTCTGAAAGTGCCACACCAGAAAGTGATAGTGCTACCCATATTAAATTCTCAAAACGTGATGAGGACGGCAAAGAGTTAGCTGGTGCAACTATGGAGTTGCGTGATTCATCTGGTAAAACTATTAGTACATGGATTTCAGATGGACAAGTGAAAGATTTCTACCTGTATCCAGGAAAATATACATTTGTCGAAACCGCAGCACCAGACGGTTATGAGGTAGCAACTGCTATTACCTTTACAGTTAATGAGCAAGGTCAGGTTACTGTAAATGGCAGCAGCGGCCTGGTGCCGCGCGGCAGCCATATGGGCCTGGGCTCCCACCACCACCACCACCACTGAGATCCGGCTGCTAACAAAGCCCGAAAGGAAGCTGAGTTGGCTGCTGCCACCGCTGAGCAATAACTAGCATAACCCCTTGGGGCCTCTAAACGGGTCTTGAGGGGTTTTTTGCTGAAAGGAGGAACTATATCCGGATTGGCGAATGGGACGCGCCCTGTAGCGGCGCATTAAGCGCGGCGGGTGTGGTGGTTACGCGCAGCGTGACCGCTACACTTGCCAGCGCCCTAGCGCCCGCTCCTTTCGCTTTCTTCCCTTCCTTTCTCGCCACGTTCGCCGGCTTTCCCCGTCAAGCTCTAAATCGGGGGCTCCCTTTAGGGTTCCGATTTAGTGCTTTACGGCACCTCGACCCCAAAAAACTTGATTAGGGTGATGGTTCACGTAGTGGGCCATCGCCCTGATAGACGGTTTTTCGCCCTTTGACGTTGGAGTCCACGTTCTTTAATAGTGGACTCTTGTTCCAAACTGGAACAACACTCAACCCTATCTCGGTCTATTCTTTTGATTTATAAGGGATTTTGCCGATTTCGGCCTATTGGTTAAAAAATGAGCTGATTTAACAAAAATTTAACGCGAATTTTAACAAAATATTAACGTTTACAATTTCAGGTGGCACTTTTCGGGGAAATGTGCGCGGAACCCCTATTTGTTTATTTTTCTAAATACATTCAAATATGTATCCGCTCATGAATTAATTCTTAGAAAAACTCATCGAGCATCAAATGAAACTGCAATTTATTCATATCAGGATTATCAATACCATATTTTTGAAAAAGCCGTTTCTGTAATGAAGGAGAAAACTCACCGAGGCAGTTCCATAGGATGGCAAGATCCTGGTATCGGTCTGCGATTCCGACTCGTCCAACATCAATACAACCTATTAATTTCCCCTCGTCAAAAATAAGGTTATCAAGTGAGAAATCACCATGAGTGACGACTGAATCCGGTGAGAATGGCAAAAGTTTATGCATTTCTTTCCAGACTTGTTCAACAGGCCAGCCATTACGCTCGTCATCAAAATCACTCGCATCAACCAAACCGTTATTCATTCGTGATTGCGCCTGAGCGAGACGAAATACGCGATCGCTGTTAAAAGGACAATTACAAACAGGAATCGAATGCAACCGGCGCAGGAACACTGCCAGCGCATCAACAATATTT |
| SpyT2 insert | GTTCCAACAATCGTTATGGTTGACGCTTACAAACGTTACAAG |
| SnpCatcher | AAGCCGCTGCGTGGTGCCGTGTTTAGCCTGCAGAAACAGCATCCCGACTATCCCGATATCTATGGCGCGATTGATCAGAATGGGACCTATCAAAATGTGCGTACCGGCGAAGATGGTAAACTGACCTTTAAGAATCTGAGCGATGGCAAATATCGCCTGTTTGAAAATAGCGAACCCGCTGGCTATAAACCGGTGCAGAATAAGCCGATTGTGGCGTTTCAGATTGTGAATGGCGAAGTGCGTGATGTGACCAGCATTGTGCCGCAGGATATTCCGGCTACATATGAATTTACCAACGGTAAACATTATATCACCAATGAACCGATACCGCCGAAA |
| SnpTag | AAATTAGGCGATATTGAGTTCATTAAGGTGAACAAA |
| I19 | GATCTGGGTAAAAAACTGCTGGAAGCAGCACGTGCAGGTCAGGATGATGAAGTTCGTATTCTGATGGCAAATGGTGCAGATGTTAATGCCGATGATAATAAAGGTGATACACCGCTGCATCTGGCAGCAAGCTTTGGTCATCTGGAAATTGTTGAAGTGCTGCTGAAAAACGGTGCCGATGTGAACGCAGATGATTATTTTGGTGATACCCCTCTGCACTTAGCAGCATGGTCAGGCCATTTAGAAATCGTGGAAGTGTTACTGAAATATGGTGCGGACGTGAATGCACAGGATCAGCGTGGTTTTACCCCGTTACACCTGGCAGCCATTGCAGGCCACCTGGAAATAGTAGAGGTTCTGCTTAAATATGGCGCTGATGTTAACGCCCAGGATAAATTTGGTAAAACCGCCTTTGATATCAGCATCGATAATGGCAATGAAGATCTGGCAGAAATTCTGCAG |
| YMB | GGTAGCAGCGTGAGCAGCGTTCCGACCAAACTGGAAGTTGTTGCAGCAACCCCGACCAGCCTGCTGATTAGCTGGGATGCAAGCAGCAGCAGCGTTAGCTATTATCGTATTACCTATGGTGAAACCGGTGGTAATAGTCCGGTTCAAGAATTTACCGTTCCGGGTAGCAGCAGTACCGCAACCATTAGCGGTCTGAGTCCGGGTGTTGATTATACCATTACCGTTTATGCCTACTACAGCTATTACGATCTGTATTATAGCTATTCACCGAGCAGCATTAACTATCGTACC |
| GFP | ATGAGTAAAGGAGAAGAACTTTTCACTGGAGTTGTCCCAATTCTTGTTGAATTAGATGGTGATGTTAATGGGCACAAATTTTCTGTCAGTGGAGAGGGTGAAGGTGATGCAACATACGGAAAACTTACCCTTAAATTTATTTGCACTACTGGAAAACTACCTGTTCCTTGGCCAACACTTGTCACTACTCTGACGTATGGTGTTCAATGCTTTTCCCGTTATCCGGATCACATGAAACGGCATGACTTTTTCAAGAGTGCCATGCCCGAAGGTTATGTACAGGAACGCACTATATTCTTCAAAGATGACGGGAACTACAAGACGCGTGCTGAAGTCAAGTTTGAAGGTGATACCCTTGTTAATCGTATCGAGTTAAAAGGTATTGATTTTAAAGAAGATGGAAACATTCTCGGACACAAACTCGAGTACAACTATAACTCACACAATGTATACATCACGGCAGACAAACAAAAGAATGGAATCAAAGCTAACTTCAAAATTCGCCACAACATTGAAGATGGTTCCGTTCAACTAGCAGACCATTATCAACAAAATACTCCAATTGGCGATGGCCCTGTCCTTTTACCAGACAACCATTACCTGTCGACACAATCTGCCCTTTCGAAAGATCCCAACGAAAAGCGTGACCACATGGTCCTTCTTGAGTTTGTAACTGCTGCTGGGATTACACATGGCATGGATGAGCTCTACAAA |
| Protein G | CTGCCGAAAACCGATACCTATAAACTGATTCTGAATGGCAAAACCCTGAAAGGTGAAACCACCACCGAAGCAGTTGATGCAGCAACCGCAGAAAAAGTCTTTAAACAGTATGCCAATGATAATGGCGTTGATGGTGAATGGACCTATGATGATGCAACCAAAACCTTTACCGTTACCGAAAAACCGGAAGTTATTGATGCAAGCGAACTGACACCGGCAGTTACCACATATAAACTGGTGATTAATGGTAAAACGCTGAAGGGCGAGACAACAACCGAAGCCGTGGACGCAGCCACAGCCGAGAAAGTTTTCAAACAATATGCAAACGACAACGGTGTGGATGGCGAGTGGACATATGACGACGCCACAAAAACATTTACAGTGACAGAGAAACCTGAAGTGATCGATGCCTCAGAACTGACCCCTGCCGTGACAACCTACAAACTGGTTATCAACGGAAAAACCTTAAAAGGCGAAACAACGACAAAAGCCGTTGATGCCGAAACAGCGGAGAAAGCATTTAAGCAGTACGCGAACGATAACGGCGTGGACGGTGTTTGGACATACGATGATGCGACTAAGACGTTTACGGTGACCGAA |

| PCR primer | Sequence |
| --- | --- |
| F Module GA | CATATGGGCCTGGGCTCCCA |
| R Module GA | ACTTTCTGGTGTGGCACTTTCAG |

**Table S2.3.** DNA sequences for all PCR primers used.

**2.4 Molecular biology of module construction**

DNA amplification used the proof-reading polymerases Q5 DNA polymerase (NEB) or Phusion (ThermoFisher). Cloning of the modular building blocks used the primers ‘F Module GA’ and ‘R Module GA’ (**Table S2.3**) to amplify the vector backbone (typically pET28a SNAP-SpyC, **Table S2.2**), followed by DpnI digestion (FD-DpnI; ThermoFisher) to remove template. Inserts were ordered as synthetic oligonucleotides (Sigma) or genestrings (GeneArt) with homology regions to the vector for Gibson assembly. PhoCl was cloned from the plasmid pBAD/HisB-PhoCl-MBP. We thank R.E. Campbell’s group (University of Alberta) for making this plasmid available through AddGene. ^1^

**2.5 Mammalian cell culture**

HeLa cells were continuously cultured in T-75 flasks (Nunc 156499) using DMEM (high glucose, GlutaMAX^TM^; Gibco 61965-026) supplemented with FBS (10 % v/v; Gibco 10500064) and passaged by washing with PBS (Merck D8537) before incubating 10 min in Accutase (Merck A6964) then resuspending in excess medium and reseeding. All incubations are at 37 C, 7 % CO_2_, ambient O_2_ (unless otherwise stated). When cells were seeded for assays, an additional aliquot was reseeded into culture for at least 2 days before testing for mycoplasma contamination using MycoAlert (Lonza LT07-318) according to the manufacturer’s instructions. All cultures used in this work tested negative.

**2.6 Fluorescent, enzymatic and phenotypic assays**

**2.6.1 Flow cytometry**

Flow cytometric analysis of beads was carried out on an Attune NxT instrument. Data was taken as the median value of a desired channel, and where triplicates are shown is the mean of the three median values.

**2.6.2 Apopotosis assays**

For the apoptosis assays in Figure 6, a suspension of freshly detached HeLa cells was pelleted 5 min at 500 *g* and resuspended in fresh medium (DMEM) before seeding at a density of 10^4^ cells per well of a 96-well flat-bottom tissue culture plate (CellStar 655180) and incubating overnight. Treatment media were prepared by normalising constructs to 115 nM effective scFv concentration in medium, then serially diluting four-fold in medium for a total of 8 treatments per construct. All treatment media were supplemented with 3.3 µg/mL cycloheximide for sensitisation (Dr. Sylwia Mankowska, personal communication). Cell medium was aspirated and replaced with 50 µL treatment media in a semi- and pseudo-randomised plate layout: constructs were assigned a random plate column, and the dosage order was randomised within each of these columns; both randomisations used the Python language’s standard ‘random’ package function, shuffle^2,3^, and all replicates were independently randomised. Plates were incubated for two hours before removing media, washing twice with 100 µL EDTA (Merck E8008), incubating 10 min in 50 µL Accutase then neutralising with 150 µL medium. The supernatant, washes, and cell suspension were all pooled into a deepwell plate (Greiner 780270), pelleted 5 min at 1000 g and the supernatant removed before resuspending in 40 µL NucView 488 staining solution (Biotium 30029) in PBS, prepared according to the manufacturer’s instructions. After 15 minutes’ incubation at room temperature, 160 µL PBS was added before transferring to a non-treated flat-bottom plate (Thermo Scientific 260836) and analysing immediately on an Attune NxT. Voltages: FSC 30, SSC 220, BluFL1 240. 100µL acquisition volume, 180 µL sample volume, 2 mixing cycles, 1 rinse cycle, 200 µL/min flow rate, up to 10000 gated events (cells).

**2.6.3 Bacteriolytic assays**

Static overnight cultures of induced *E coli* cells (BL21 DE3, expressing GFP-SpyTag) were incubated with the indicated carbenicillin concentrations (+/- PBS dilution) in 200 µl volumes in individual wells of a 96-well deep-well V-bottomed plate. In parallel an aliquot (the same volume as for static culture conditions) of the overnight static cultures was centrifuged at 7000*g* for 5 minutes and the cell pellet resuspended in an equal volume of fresh media, these samples were then handled as both the static and PBS dilution samples were. Samples were then incubated with shaking (200 rpm, 37 ^o^C, 90 minutes) to induce bacteriolysis. The cell debris/cells were then pelleted at 4000 *g* for 10 minutes, and the supernatants (150 µl) transferred to a new plate for capture of released GFP-SpyTag with 50,000 20 µm, 50 µM BG SNAP-SpyCatcher functionalised beads for 60 minutes with shaking. After 60 minutes the beads were pelleted at 4000 *g* for 5 minutes, the supernatant removed and 1 ml wash buffer added. This process was repeated twice. Subsequently beads were analysed for GFP-SPyTag capture, and therefore the degree of induced bacteriolysis, by flow cytometry (Figure 5d).

**2.6.4 Analysis of P91 enzymatic activity on bead**

The described (Figure 5b) number of beads functionalised with P91-SpyTag, were incubated with 50 µM fluorogenic phosphotriester for 12 hours (Kind gift of J.D.F. Schnettler-Fernandez). An increase in fluorescence in the green channel due to fluorescein release was followed in a Tecan spectrophotometer (Tecan Infinite 200 Pro). Catalytic activity was calculated by taking the first 90 minutes of data collected and calculating the slope.

(2)	Van Rossum, G., 2020. *The Python Library Reference, release 3.8.2*, Python Software Foundation.

(3)	https://github.com/fhlab/Fryer-et-al-2021

(4)	Diamante, L.; Gatti-Lafranconi, P.; Schaerli, Y.; Hollfelder, F., In vitro affinity screening of protein and peptide binders by megavalent bead surface display. *Protein*

*Eng Des Sel* **2013**, 26 (10), 713-24.

(5)	Daramola, O.; Stevenson, J.; Dean, G.; Hatton, D.; Pettman, G.; Holmes, W.; Field, R., A high-yielding CHO transient system: Coexpression of genes encoding EBNA-1 and GS enhances transient protein expression. *Biotechnology Progress* **2014** 30, 1, 132-141.
